# Double Trouble: Multiple infections and the coevolution of virulence and resistance in nested host-parasite populations

**DOI:** 10.1101/2024.07.17.603867

**Authors:** Julien D. Lombard, François Massol, Sébastien Lion

## Abstract

Parasite competition is a key factor driving the epidemiology and evolution of parasites, and is expected to modulate the selective pressures acting upon the host, and to alter its response to infection. Using a nested co-evolutionary model where epidemiological traits (transmission, virulence and recovery) are derived from within-host interactions between parasite strains, we analyse using adaptive dynamics methodology the joint evolution of parasite virulence and host resistance. We compare the predicted coevolutionarily stable states under different competition regimes, including single infections (preemption), superinfection (strain dominance) and coinfections (transient strain coexistence). We find that coinfections select for lower virulence compared to superinfection, while the opposite trend is observed for host resistance. The local coexistence of parasites in coinfected hosts creates a kin selection effect that reduces both parasite virulence and host response. We show that the magnitude of this effect depends on the ecological context, and notably the degree of coupling between hosts. Our work contributes to building the gap between metapopulation ecology and epidemiology.

## Introduction

The study of host-parasite coevolution is of major interest in a wide variety of scientific fields including agronomy, conservation biology, and human or animal health. The antagonistic interaction between parasites and their hosts leads to complex eco-evolutionary feedback loops. Assessing how evolutionary forces shape and are shaped by environmental or physiological mechanisms is thus a difficult task. Since the seminal work of May and Anderson (1979), there have been numerous attempts at drawing a mathematical picture of the evolution of host-parasite interactions. In particular, a strong effort has been dedicated to studying coevolution in attack-defense traits. There is increasing theoretical and empirical support showing that the coevolution of parasite virulence and host resistance are crucial determinants of the ecology and evolution of infectious diseases (Restif and Koella, 2003; Best et al., 2009; Carval and Ferriere, 2010; Kada and Lion, 2015 for the theory; Webster et al., 2004; Lefèvre et al., 2007; Frickel et al., 2016 for the empirical part). An important feature of infections is that hosts are often infected with multiple parasite strains that (at least transiently) coexist in the host (*e*.*g*. Susi et al., 2015). Multiple infections, and more specifically interactions between different genotypes of parasites within the host, are known to shape the ecological outcome of infection (Mideo, 2009; Day et al., 2011; Sofonea et al., 2015), reverberating at the between-host scale. Competition among parasites within the host, together with the subsequent host physiological or immune response, generates selective pressures (De Roode et al., 2005). These pressures may act in opposite directions across scales, favoring for example increased virulence at the within-host scale but reduced virulence at the between-host scale (reviewed in Mideo et al., 2008; Cressler et al., 2016). However, such complex eco-evolutionary dynamics remain poorly accounted for in theoretical studies, especially in a coevolutionary framework.

Amongst epidemiological models analyzing the evolution of virulence, most –if not all– rely on phenomenological assumptions to capture the outcome of within-host competition dynamics. From our literature survey, three main approaches are traditionally used. The classic approach is to neglect multiple infections altogether: infected hosts are then assumed to be immune to reinfection by other strains (May and Anderson, 1979; Restif and Koella, 2003; Day et al., 2011; Best et al., 2012). Thus, competition is assumed to be ruled exclusively by the colonization of new susceptible individuals at the between-host scale. A second approach relies on the superinfection assumption, in which the competition between two coinfecting strains is instantaneously resolved by the exclusion of the less competitive one (Kada and Lion, 2015; Nowak and May, 1994). The underlying assumption is that parasite strains respect a strict competitive hierarchy. While this hypothesis may hold when competitive differences are strong, it is, however, unlikely for multiple infections by genetically close strains. A third approach is to explicitly allow strains to coexist within a host (Sofonea et al., 2015; May and Nowak, 1995; Mosquera and Adler, 1998). Alizon and van Baalen (2008) presented a multiple-infection model in which coinfections emerged as an outcome of within-host dynamics. Their model include mixed regimes of competition because a strain with a competitive advantage will slowly displace its competitor, but might not succeed to fully replace the other before the infection ends. One of their main findings was that allowing for transient coexistence of competing strains makes the evolutionary branching of virulence more likely when coinfections are frequent. This result is a direct consequence of the incorporation of multiple infections, which creates an heterogeneity in the types of hosts that can be infected.

Multiple infections in epidemiology share conceptual similarities with metapopulation ecology, because a host can be seen as a local habitat in which several parasite strains interact. Competition for new susceptibles in single-infection models acts in a very similar way to preemptive competition in metapopulations models (*i*.*e* the resident status of a strain prevents further competitive replacement by another). Superinfection models, on the other hand, describe competitive rules that are very similar to the ones defining the competition-colonization trade-off model (Hastings, 1980; Tilman, 1994) from Levins (1969) seminal formalism. In such cases, better competitors instantaneoulsy displace less competitive species, such that competition acts through strict dominance. Allowing for intermediate cases between complete dominance or preemption in the description of species competition has been shown, for example, to alter the likelihood of coexistence at the regional scale (Calcagno et al., 2006). In particular, the coexistence of species or strains at the regional scale should be favored when the fitness is not completely determined by the colonization process (*i*.*e*. the transmission to suceptible hosts in an epidemiological context). The integration of mixed effect of dominance and preemption in strain competition during multiple infections thus appears to be a relevant feature to include in coevolutionary studies.

The occurrence of multiple infections can directly influence how hosts respond to parasites. Reciprocally, host resistance can alter how selection acts on parasite virulence when parasites compete within the host. When acting as an avoidance mechanism, resistance reduces transmission rates and thus the likelihood of multiple infections (Boots and Haraguchi, 1999), which in turn can affect the strength of dominance competition. When acting as a clearance mechanism, the host resistance reduces the infectious period (van Baalen, 1998), which should increase the strength of preemptive competition, because less time is allowed for the strains to compete. The relative contribution of preemption and dominance effects in strain competition is therefore expected to affect the importance of a given competition scale. There is thus a need to incorporate multiple infections and detailed within-host interactions processes in a coevolutionary framework. However, despite notable theoretical advances in the study of host-parasite evolution, such mechanisms remain poorly included, especially in a coevolutionary perspective.

Here, we build on the existing theory of metapopulation ecology and develop a nested model to study the coevolution of host resistance and parasite virulence. From individual-based traits, we define a within-host model including parasite growth and host-induced parasite suppression. Then, using a time-scale separation argument, we derive from the within-host equilibrium the main processes (transmission, virulence, recovery) of the between-host epidemiological model. Our results are derived assuming trade-offs between parasite replication and virulence, and host resistance and fecundity. Using the now classic toolbox of adaptive dynamics (Geritz et al., 1998), we first provide an overview of the evolutionary outcomes resulting from the evolution of either the host or the parasite. Then we consider the coevolution of both partners, and determine how strain competition and host response drive selection of host resistance and parasite virulence.

## Epidemiological dynamics

### SIS epidemiological model

Our starting point is the classic susceptible-infected-susceptible (SIS) model (Kermack and McKendrick, 1927). Because, from the parasite’s point of view, the host population can be viewed as a well-mixed metapopulation of fully connected patches, we reframe the SIS dynamics as a metatpopulation model, and track the dynamics of the fractions of patches in each state. Here, each patch can be either empty (0) or occupied by a single susceptible (*S*) or infected host (*I*). Assuming that density dependence acts on natality through the occupancy of the host metapopulation (*i*.*e*. no further birth event can occur once all patches are occupied), the demographic and epidemiological dynamics of the host are given as:

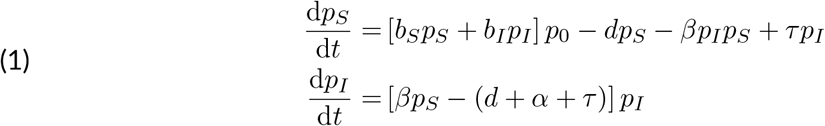

where *p*_*x*_ is the fraction of patches in state *x*. Note that our model reduces to two equations because the patch frequencies sum to one (*p*_*S*_ + *p*_*I*_ + *p*_0_ = 1). Susceptible (resp. infected) hosts reproduce at a rate *b*_*S*_ (resp. *b*_*I*_), and give birth to susceptible newborns, which can only develop into adults if they find an empty patch, with probability *p*_0_. Both susceptible and infected hosts have a background mortality rate *d*. Infected hosts have an additional mortality rate (virulence) *α*, and can recover at a rate *τ*. Finally, disease transmission occurs at a rate *β*.

Provided that *b*_*s*_ *> d*, the model has two non-trivial equilibria: a disease-free equilibrium in the form 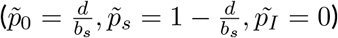, and an endemic equilibrium 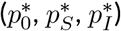. For the endemic equilibrium to exist, the basic reproduction number *R*_0_, that is the expected number of secondary infections produced by a single infected host in an otherwise uninfected population (Diekmann et al., 1990), must be greater than 1, that is

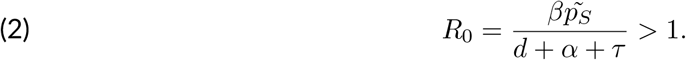

Note that, in our metapopulation formalism, *R*_0_ is equivalent to the number of successful dispersers produced by an initial parasite infecting a host in an otherwise uninfected host population and in the absence of evolution, which is typically noted *R*_*m*_ in the metapopulation literature (Metz and Gyllenberg, 2001; Ajar, 2003; Massol et al., 2009).

### Within-host model

Following Alizon and van Baalen (2005), we derive the epidemic traits from within-host interactions. We introduce a model of within-host parasite growth which tracks the dynamics of the within-host parasite density, noted *X*:

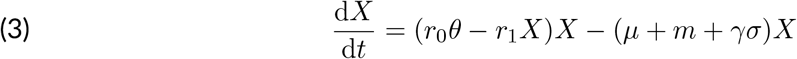

Parasites replicate at a rate *r*_0_*θ*, where *r*_0_ is the maximum replication rate and *θ* is the parasite investement in host exploitation, which takes values between 0 and 1. We assume that within-host replication is density-dependent, with strength *r*_1_. Parasite emigrate from their host at a rate *m* and die at a rate *µ* + *γσ*, where *µ* refers to background parasite mortality and *γσ* is the host resistance level. *γ* is a theoretical maximum for resistance, and *σ* is the host investment in resistance, which takes values between 0 and 1. The within-host dynamics then have two possible outcomes: either the parasite becomes extinct 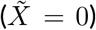, or its density stabilises at an equilibrium parasite load, *X*^∗^, given by:

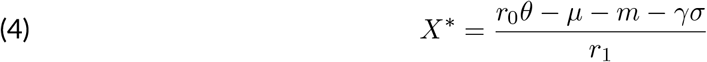

Thus, higher within-host replication leads to higher within-host loads. The next step in our modelling approach is to make epidemiological parameters depend on the within-host load, leading to explicit trade-offs between transmission, virulence, and recovery (Frank, 1996).

### Bridging the scales

System (1) describes a well-mixed population. However, from the parasite perspective, the host population can be seen as a set of discrete pacthes linked through the dispersal of parasite propagules. Our SIS model can therefore describe the colonization-extinction dynamics of local parasite populations (Levins, 1969), the metapopulation dynamics of which being given by eqn.(1). Thus transmission can be seen as the rate at which new host pacthes are successfully colonized, recovery as a local extinction of the parasite population where the patch remains available, and the host death as a particular extinction event where the patch remains temporarily unavailable (until a new host is born). To ensure that the coupling between system (1) and eqn. (3) remains consistent, we assume that within-host dynamics is fast compared to the processes that put an end to infection (*i*.*e*. host mortality, recovery and virulence). This requires that the rate at which eq.(3) shifts from equilibrium 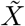 to *X*^∗^ is much greater than the rate at which hosts leave the infected class, which mathematically translates into :

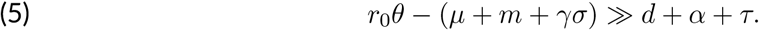

### Epidemiological traits

To make further progress, we assume that epidemiological parameters depend on the within-host equilibrium load *X*^∗^. We assume that transmission is a linear increasing function of the equilibrium load, so that

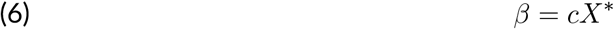

where *c* = *m*(1 − *ρ*) is the rate of successful dispersal of parasite propagules, *m* the rate at which propagules leave their host, and *ρ* a cost that represents the fraction of emitted propagules lost during dispersal. Parasite transmission is thus equivalent to the rate of successful dispersal events. Note that we implicitly consider, for simplicity, that the time spent by propagules during transmission is negligible, so that emitted propagules either immediately infect an other host, or die trying.

For virulence and recovery, we consider non-linear trade-off functions

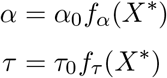

where *f*_*α*_ (resp. *f*_*τ*_) are increasing (resp. decreasing) functions of the within-host parasite load and *α*_0_ (resp. *τ*_0_) is a theoretical maximum for the virulence (resp. recovery). The functions *α* and *τ* also depend on a shape parameter used to vary the shape of the trade-off function (from concave to convex). A complete description of the functions used is given in appendix C. In addition, a table summarising parameter notations and interpretations is provided in appendix A.

Because the within-host density *X*^∗^ is affected by the host’s investment into defence *σ* and the parasite’s investment into host exploitation *θ*, the epidemiological parameters implicitly depend on these two traits. In particular, because *X*^∗^ is an increasing function of *θ*, both transmission and virulence are assumed to be increasing functions of the within-host replication rate, which has received strong empirical support (Acevedo et al., 2019). As a result, as in the classic virulence-transmission trade-off hypothesis, an increase in transmission can only be bought at the expense of a reduced infectious period (Alizon et al., 2009).

In addition, we assume that host defense is costly and leads to a reduced fecundity of infected hosts, that is :

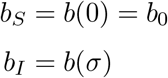

where *b*(*σ*) is a decreasing function of *σ*. This implies that there are no constitutive costs of resistance, because only infected hosts pay a cost.

Note that resistance in our model acts in two ways: as a mechanism that causes epidemic mitigation (*i*.*e*. causing a reduction in disease transmission), and as a clearance mechanism (*i*.*e*. causing an increase in host recovery, and/or a decrease in virulence).

## Evolution of host resistance

In this section, we assume that parasite investment in replication *θ* is fixed, and only consider the evolution of host resistance, *σ*.

### Host and parasite coexistence

Following the infection of a susceptible host, the within-host parasite density will converge to a positive equilibrium *X*^∗^ if and only if

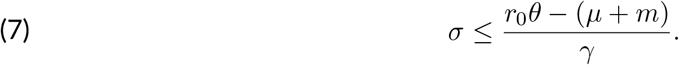

Because *σ* takes values between 0 and 1, this implies that the host always clears the disease when *r*_0_*θ < µ* + *m* and that the host always becomes infected with parasite load *X*^∗^ when *θ >* (*γ*+*µ*+*m*)*/r*_0_. For simplicity, we will now restrict our analysis to scenarios where the host is never able to eradicate the disease, irrespective of its level of resistance. However, an overview of cases where the host evolution can drive the parasite towards extinction is presented in appendix B.

In addition, an endemic equilibrium exists at the population level if *R*_0_ *>* 1. Note that if *R*_0_ *>* 1, this implies that the inequality (7) is also verified.

It also follows that for a given level of host resistance *σ*, a parasite that does not sufficiently invest in virulence cannot establish a within-host viable population and therefore has its *R*_0_ *<* 1, leading to its global extinction (figure 2a).

**Figure 1.**
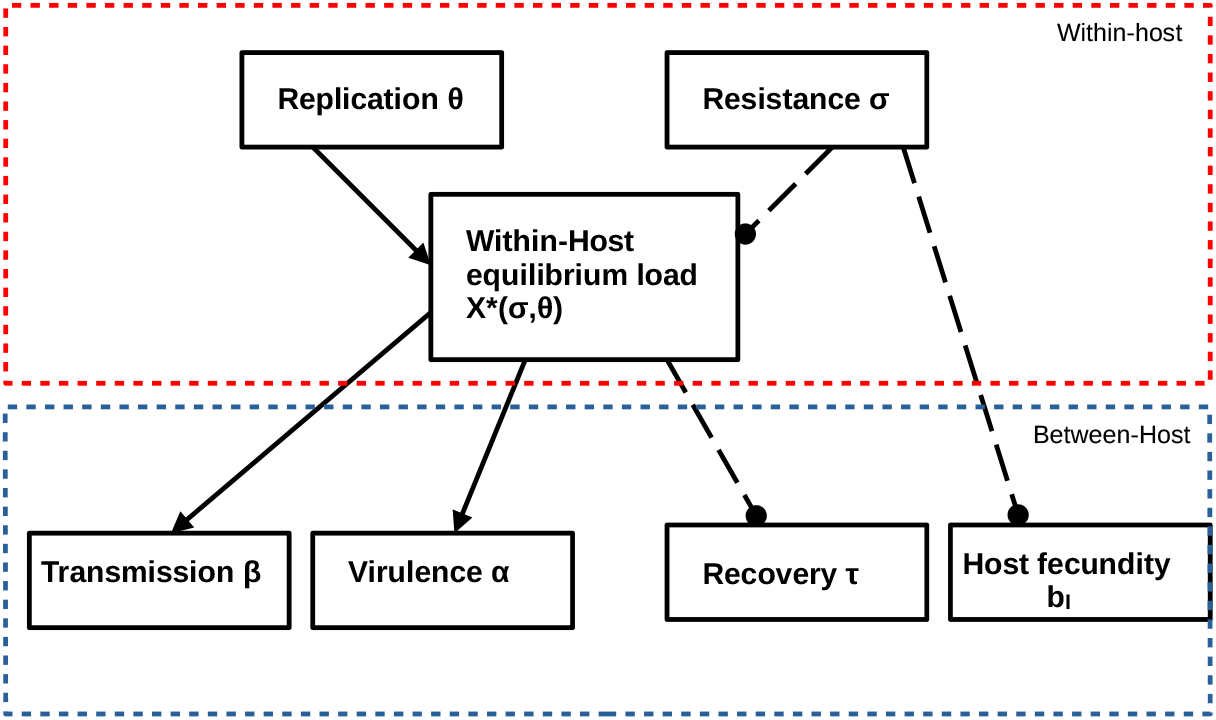
Schematic view of interactions between resistance and exploitation traits, as well as their consequences on equilibrium density *X*^∗^ and associated epidemiological parameters. Solid arrows refers to up-regulation links (when the origin quantity increases, it induces an increase in the terminal quantity). Dashed lines with black circles refers to inhibition links (when the origin quantity increases, it induces a decrease in the terminal quantity). The red square delimits processes acting at the within-host level, while the blue square delimits between-host processes.

**Figure 2.**
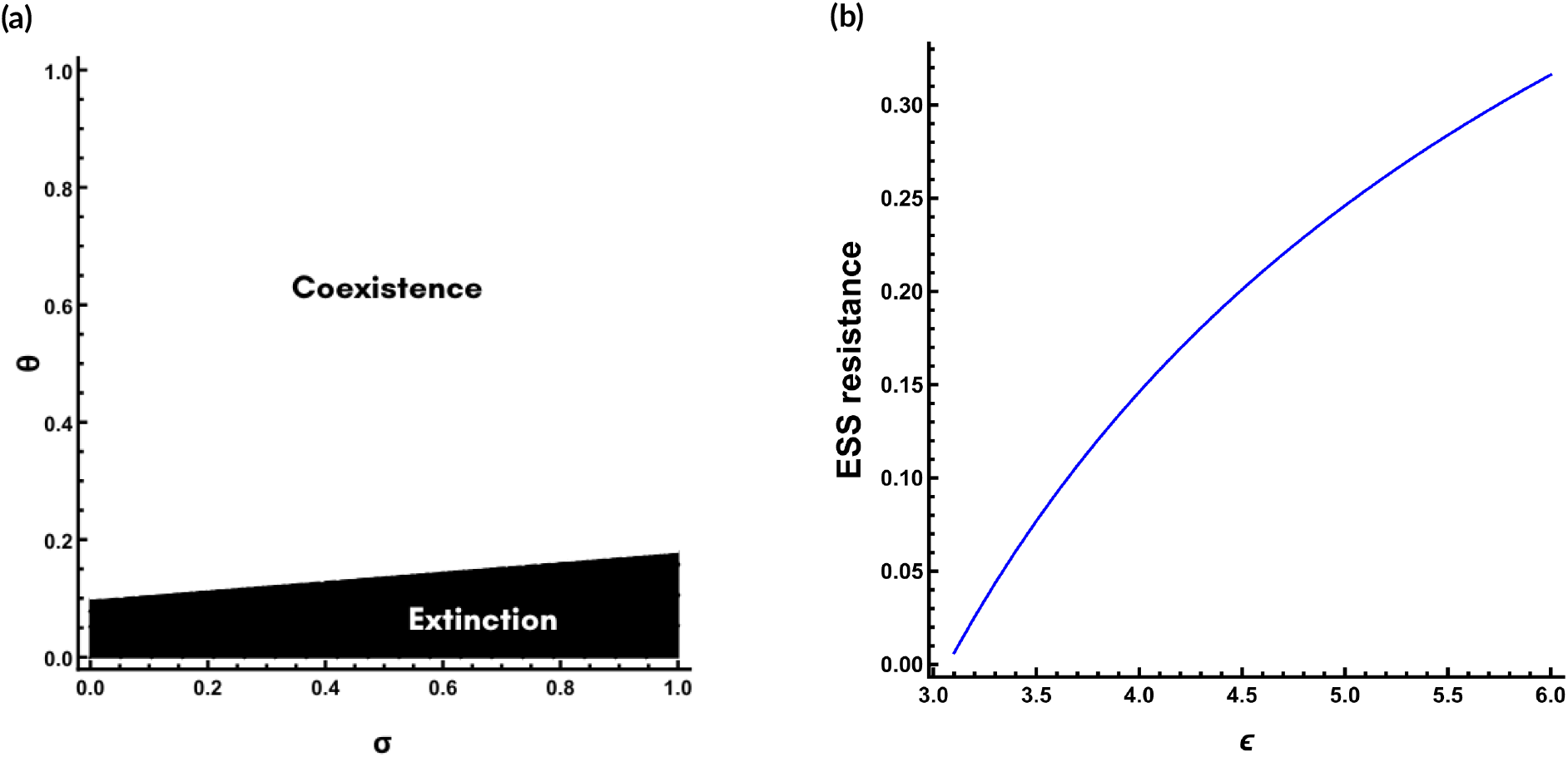
(a) Phase diagram describing the parasite domain of existence. White areas represent the cases where the hosts and parasite always coexist, while black areas represent the set of cases where the host can drive its parasite towards extinction. (b) Existing and biologically relevant CSS strategies of host investment in resistance as a function of the shape of the trade-off function between host resistance and fecundity, *ϵ* (concave when *ϵ >* 1, linear when *ϵ* = 1, convex when *ϵ <* 1). Other parameter values were set to :*b*_0_ = 0.8, *µ* = 0.5, *α*_0_ = 1, *r*_0_ = 10, *θ* = 0.8, *r*_1_ = 0.04, *ρ* = 0.9, *γ*_0_ = 1, *d*_0_ = 0.05, *τ*_0_ = 1, *δ* = 3, *κ* = 1, *d* = 0.5.

### Invasion Fitness

We consider the fate of a rare mutant host with investment *σ*_*m*_ invading a resident host population with investment *σ* at its endemic equilibrium. We define the invasion fitness as the initial growth rate of the mutant in the equilibrium resident population (Geritz et al., 1998). For structured populations, a proxy for the invasion growth rate is obtained using the next-generation theorem (Diekmann et al., 1990; Hurford et al., 2010; Otto and Day, 2007). Denoting quantities that depends on the mutant trait by a subscript *m*, this yields (see appendix D for the complete derivation) :

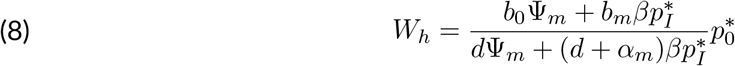

where Ψ_*m*_ = *d* + *α*_*m*_ + *τ*_*m*_, such that 1*/*Ψ_*m*_ is the average infectious period of a mutant host.

Eqn.(8) is similar to eqn. (A.7) in Kada and Lion (2015) or eqn. (6) in Restif and Koella (2003), with some noticeable differences. First, we assume that the mutation affects both virulence and recovery. Second, the birth rate of susceptible individuals does not depend on the investment in resistance, as we assume the costs of resistance only come from mounting an immune response. It is then straightforward to see that in the absence of the parasite 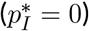, resistance becomes a neutral trait (as 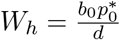, which is independent of the mutant trait). As only infected individuals pay the costs of resistance, parasite presence is needed for resistance to be under selection.

### Singular strategies

Following the standard adaptive dynamics methodology, the host’s evolutionary singularities are obtained by calculating the zeros of the host selection gradient 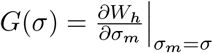. Evolutionary stability is then determined using the sign of the second derivatives of the fitness function. A strategy is called *evolutionarily stable* (ESS) if the second partial derivative of the fitness function with respect to the mutant trait is negative (Geritz et al., 1998). In addition, if the derivative of the fitness gradient *G*(*σ*) evaluated at the singular strategy is negative, the strategy is said to be *convergence stable*, which ensures that it is a locally attracting point. A strategy that is both evolutionarily stable and convergence stable is thus an endpoint of the evolutionary dynamics and is termed as a *continuously stable strategy* (CSS) (Eshel, 1983). A strategy that is not convergence stable is called an evolutionary repellor.

Within the range of functions and parameters explored, CSSs were found only for sufficiently concave forms (*i*.*e. ϵ* > 1), and not for linear or convex trade-offs, which is in line with patterns already observed by Best et al. (2009) and Boots and Haraguchi (1999). Biologically, this means that an evolutionarily stable investment in resistance occurs only when the subsequent physiological costs are accelerating. When the costs are moderately decelerating, the evolutionarily singular point is a repellor that lies outside of the parameter region that is biologically relevant. In such a case, selection drives resistance towards zero investment. When the costs are strongly decelerating, the repellor is displaced in the biologically relevant parameters region, and selection drives resistance towards an all-or-nothing strategy, depending on the initial conditions. Pairwise invasibility plots for decelerating costs are presented in appendix A.

When a CSS level of investment exists, it increases with the trade-off concavity (Fig.2b). In such a case, higher investment levels can be reached without paying an excessive cost, because of the increased concavity of *b*(*σ*). Unlike previously cited studies, we found no evidence for evolutionary branching under the parameter ranges explored, in contrast to Best et al. (2009) and Boots and Haraguchi (1999). This may reflect the specific trade-off shapes considered for virulence and recovery, or the feedback structure induced by immune costs that are paid only during disease carriage, both of which may constrain the invasion fitness landscape to directional selection.

## Parasite evolution

We now turn to the evolutionary dynamics of parasite investment, under the assumption that host resistance is not subject to selection. Due to the metapopulation structure of the parasite, it is not possible anymore to define fitness as the initial growth rate of a mutant invader in a given host. We therefore use as a fitness measure the average lifetime population success *R*_*m*_, (Metz and Gyllenberg, 2001; Ajar, 2003; Massol et al., 2009), following the methods initially described in Jansen and Vitalis (2007) and Pillai et al. (2012).

### Competition dynamics in coinfected hosts

We consider the fate of a rare mutant parasite with trait *θ*_*m*_ and within-host equilibrium load 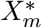 during the early stages of the invasion of a resident endemic equilibrium (with the resident *r* strain having the trait value *θ* and equilibrium load *X*^∗^). This implies that we need to consider the reinvasion dynamics of strains, which can occur when the mutant invades a host already occupied by the resident parasite, or conversely when being reinvaded by the resident while occupying the host on its own.

At a given time *t*, a host coinfected by two parasiste strains *r* and *m* can be viewed as a shared patch characterised by its frequency of mutant pathogen and its total within-host parasite load. In Appendix E, we show that the dynamics of the mutant frequency *f* (*t*) in a shared host are given by

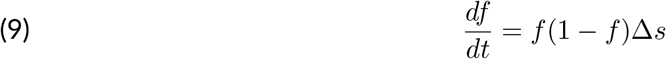

with competition acting through the differential replication of the two strains Δ*s* = *r*_0_(*θ*_*m*_ − *θ*). Because Δ*s* is independent of time, eqn. (9) can be solved in closed form. Starting from an initial mutant frequency *ϕ*, this yields:

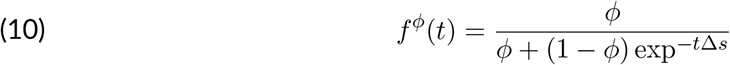

Note that, as expected, *f*^*ϕ*^(*t*) = *ϕ* when Δ*s* = 0. Depending on whether we look at the reinfection of a host infected by the resident or mutant host, we define two initial conditions of interest:

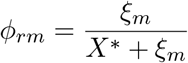, the initial mutant frequency when the mutant is the secondary infection,

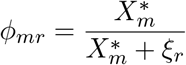, the initial mutant frequency when the resident is the secondary infection. where *ξ*_*m*_ and *ξ*_*r*_ refers to the number of invading propagules at the time of invasion (*i*.*e*. the inoculum size for the mutant and resident strains, respectively). For the following, we will use for the sake of simplicity *ξ*_*r*_ = *ξ*_*m*_ = 1 without loss of generality. Thus, a coinfected host can be in one of two states: either *rm*, in which case the host is first infected by the resident pathogen and then reinfected by the mutant, or *mr*, in which case the host is first infected by the mutant parasite and then reinfected by the resident parasite. During the coinfection, each strain will produce transmitting propagules until we reach competitive exclusion, or extinction through the death or recovery of the host. When the host death or recovery is likely to occur before exclusion, even a less competitive strain will benefit from having been the first to colonize a new host, which enables preemptive effects in competition.

Assuming that mutations have a small phenotypic effect (*i*.*e. θ*_*m*_ ≈ *θ*), it follows that the total within-host density of pathogens reaches a quasi-equilibrium on a fast time scale. In Appendix E, we show that, starting from an initial mutant frequency *ϕ* at reinvasion, we can write the total parasite density in a coinfected host *Y* ^*ϕ*^(*t*) as a weighted sum of the equilibrium densities 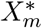 and *X*^∗^ in singly-infected hosts,

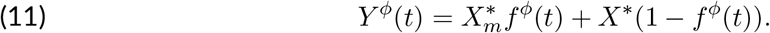

From the within-host parasite load *Y* ^*ϕ*^(*t*), we can now define the rate at which infection ceases as

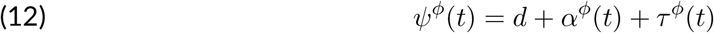

where *α*^*ϕ*^(*t*) and *τ*^*ϕ*^(*t*) are now functions of of the quasi-equilibrium load *Y* ^*ϕ*^ (eqn.11).

In order to study the dynamics of coinfection, we need to track the number of strains received by the host during the course of infection. A host can theoretically be reinfected several times without limitations. However, in order to keep the model analytically tractable, we assume that during its lifespan hosts suffer at most two infections events, and we neglect any further reinfection events. This assumption has already been used in the context of virulence evolution (van Baalen and Sabelis, 1995), or dispersal evolution (Jansen and Vitalis, 2007). In the following, it will be useful to keep track of the number of infections in a neutral model (or equivalently in a population infected by a single parasite strain). To do so, we decompose the infected compartment in system (1) according to the number of invasion experienced by a host (1 and 2), which gives

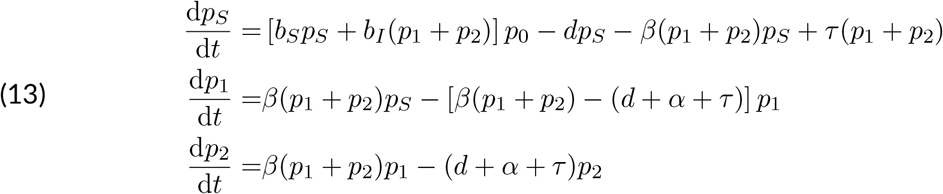

where *p*_1_ and *p*_2_ refer to the number of hosts that have received one or two parasite invasions, respectively. Using the fact that *p*_1_ + *p*_2_ = *p*_*I*_ it is straightforward to recover system (1).

### Invasion fitness

To calculate the invasion fitness, we follow the number of transmitting propagules produced by a rare focal mutant parasite landing in a given host. This number is equal to the sum of propagules produced in each of the different hosts states (susceptible or infected) in which the mutant can appear, weighted by the time spent in each of those states (Massol et al., 2009). We consider the set of fates a mutant invader can experience. If the mutant lands in a susceptible host 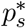, it will produce 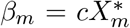 propagules per unit of time until the infection ends or the host is reinvaded. Such reinvasion will happen with probability 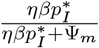, where *η* is the relative susceptibility of an infected host to reinfection. Subsequently, the mutant will produce additional propagules until competitive exclusion occurs or until the infection ends. Noting *F*^*ϕ*^(*t*) the amount of successful propagules produced after a reinvasion (either *m* → *r* or *r* → *m*), we have, using the ingredients of the previous section:

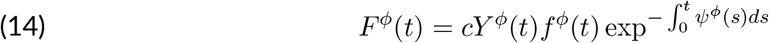

Here, *cY* ^*ϕ*^(*t*)*f*^*ϕ*^(*t*) denotes the number of mutants propagules produced at time *t*, and 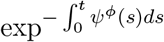 is a survival function that describes the probability that the infection has not ended by time *t*. Thus, 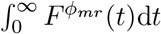 propagules are produced when the mutant is reinvaded by the resident.

Alternatively, a mutant propagule can land on an host infected by the resident pathogen, with probability 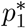 in which case it will produce 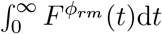 propagules over the lifespan of its host. Dropping the dependency on the resident trait for notational convenience, and denoting dependencies on the mutant by a subscript *m*, we put all these reinfection pathways together and obtain the parasite fitness function :

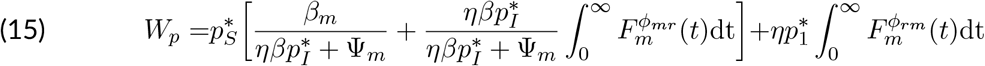

In the absence of multiple infections (*i*.*e*. when *η* = 0), parasite fitness reduces to :

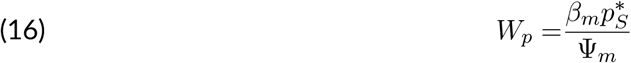

with 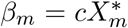. Noting 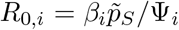 the strain-specific basic reproductive number, we have 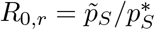, and therefore

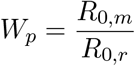

which is the classic expression for parasite fitness in the SIS model, and implies that the strain with the larger *R*_0,*i*_ value is selected (this is often called the *R*_0_ maximisation principle, see Lion and Metz, 2018).

### Fitness gradient

The complete derivation of the fitness gradient is described in appendix F. Dropping the dependency on traits to avoid notational clutter, it leads to the following expression :

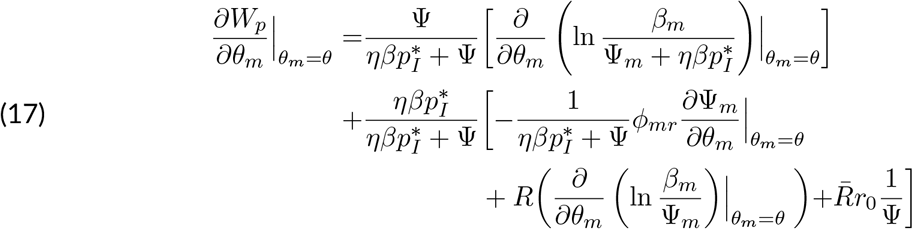

where *R* and 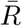 are the relatedness and unrelatedness measures in metapopulation. The relatedness *R* describes the probability to sample two identical mutants in the same host relative to the probability of sampling two identical mutants in the whole host (meta)population (Jansen and Vitalis, 2007; Rousset and Billiard, 2000). Note that this is in our case equal to the probability of interaction between two mutant indiviuals (van Baalen and Rand, 1998). Hence, the unrelatedness 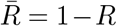 is the probability of interaction between a mutant and a resident, which can be seen as a proxy for competition intensity betwen strains. The complete derivation of the relatedness measure is described in appendix F.

To understand equation (17), it is useful to first consider what happens if we neglect multiple infections altogether. Setting *η* = 0, only the terms of the first line remain and we can rewrite the selection gradient as

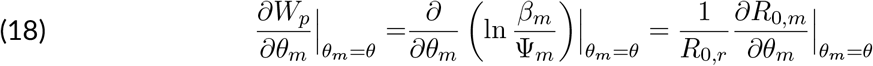

which is fully consistent with the selection gradient obtained from differentiating eq.(16). Without multiple infections, competition is only driven by preemption (i.e competition for new susceptibles), and selection pushes the parasite investment towards the highest *R*_0_. When multiple infections are possible, the interpretation of the terms on the first line remains the same, excepted that singly-infected hosts must not have been reinfected, which happens with probability 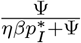, and that the average duration of a single infection is reduced by the possibility of reinfection.

The second and third lines describe how selection is affected by the occurrence of multiple infections, which happens with probability 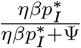. On the second line, the term between brackets corresponds to the change in the infectious period caused by the coinfection. Thus, this can be seen as the marginal effect of changing the mutant’s exploitation trait on its own clearance rate, through the change in *X*^∗^.

Finally, the terms in the third line reflect the additional selective pressures induced by the consideration of within-host competition. Interestingly, this latter component can be interpreted in terms of kin structure. When the within-host relatedness tends to 1, within-host competition occurs only among kins, and coinfected hosts tend to be similar to singly infected hosts. Hence, selection in that case should be driven by the epidemiological *R*_0_, as it would be in the case of complete preemptive competition. Conversely, when when there is no kin to compete with (*i*.*e*.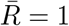), within-host selection is driven by *r*_0_ which is the marginal effect of changing the mutant exploitation on Δ*s* = *r*_0_(*θ*^′^ − *θ*), that drives the change in the within-host mutant frequency (see eqn. 9). Hence, the *r*_0_ can be understand in terms of dominance of a strain on the other. The ratio R/(1-R) therefore quantifies the balance between between-host and within-host selection (within-host selection is stronger when R=0, and between-host selection stronger when R=1), which is reminiscent of previous studies on the evolutionary consequences of multiple infections (Frank, 2012; Lion, 2013).

### The limit of pure dominance

Eqn. (18) reflects expectations about how selection should act in the limit case of pure preemptive competition. However, the limit case of pure dominance competition (*i*.*e*. close or equal to a superinfection framework) is a bit more difficult to mimic on the basis of eqn. (17). In order to adress it, preemptive effects in infected hosts must become negligible. This would correspond to infinitely long infectious periods, or strong competitive differences between strains. The first case would force crucial epidemiological parameters (virulence and recovery) to also become negligible, which would compromise the evolutionary analysis. The second case would interfere with the quasi-equilibrium assumption used to derive our fitness equation. Consequently, the derivation of pure dominance competition from eqn.(17) require additional assumptions. If one assumes that within-host competitive exclusion occurs instantaneously, restricting the number of invasions is not necessary anymore. Then, eqn.(15) drastically simplifies, and we show in appendix G that the parasite invasion fitness under the superinfection framework is:

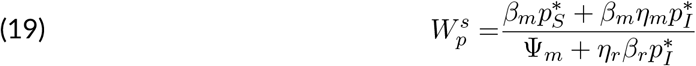

which, in the absence of the shared control of epidemiological traits, is equivalent to eq.(A3) in (Kada and Lion, 2015). Note that here, *η*_*r*_ and *η*_*m*_ are the probability that the resident (resp. mutant) strain displaces the mutant (resp. resident) upon reinvasion. In such a case, the fitness gradient becomes :

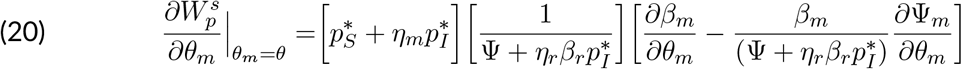

Provided that the reinvasion probabilities *η* may reflect in that case the within-host competitive ability of strains, both frameworks (*i*.*e*. superinfection or coinfection) show that the fitness of a mutant parasite is always determined by both its *R*_0_ and its within-host competitive ability. This has already been found in several theoretical works (Kada and Lion, 2015; Nowak and May, 1994; Alizon and van Baalen, 2008; van Baalen, 1998; Alizon et al., 2013) and has been shown to generally select for higher virulence. We find that the same effect occurs in our model, as competition is determined by the within-host growth rate, whose virulence is an increasing function.

### Evolutionarily singular strategies

The zeros of eqn.(17) correspond to evolutionarily singular strategies. Figure (3) shows that an evolutionarily stable investment in virulence exists only when the trade-off between replication and virulence is sufficiently convex. Otherwise, investment in virulence is always pushed towards its maximal value. Biologically speaking, this means that an intermediate evolutionarily stable investment is possible only when the costs of virulence are accelerating.

## Coevolution

We now extend our framework to coevolutionary dynamics. We first constrain the tradeoff curvatures such that an evolutionarily stable strategy exists for both species when evolving alone. The potential endpoints of host and parasite coevolutionary dynamics (co-CSS strategies) are found by studying values that cause both the host and parasite gradients to vanish. ESS-stability and convergence stability were assessed using criterions detailed in Best et al. (2009), (Appendix I).

The standalone evolutionary analysis of host resistance and parasite virulence (see eqns. (8), (17)) revealed that the force of infection and the infectious period are important drivers of within-host competition. The force of infection determines the prevalences in the host population, and thus the likelihood of multiple infections. The infectious period determines the time allowed to strain competition when multiple infections occurs. A shortened infectious period should thus enhance the part of preemption in parasite fitness (*i*.*e*. when 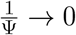 in eqn.(17)), while a long infectious period should elicit dominance competition (*i*.*e*. 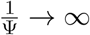 in eqn.(17)). We therefore choose to focus our coevolutionary analysis on model parameters that are most likely to affect those epidemiological features, and compare the coevolutionary outcomes under the coinfection and superinfection models.

### Dominance competition strenghtens selection towards virulent parasites and well-defended hosts

Figures (4, 6, 7) shows that the coevolutionary dynamics exhibits a single co-CSS.

Our first result is that both host resistance and parasite virulence are selected towards higher levels in the superinfection model than in the coinfection model. This can be explained by the absence of preemption in superinfections. As parasite fitness is mostly determined by dominance competition under superinfection, higher virulence and resistance levels are generally selected for. Allowing for preemptive effects by considering the local coexistence of strains (*i*.*e*. coinfections) leads to lower levels of investment for both partners. In addition, we have also shown that coinfections induce the emergence of a relatedness effect that is expected to counteract the dominance competition effects. From eqn. (17), it follows that the dominance competition factor *r*_0_ is maximal at *θ* = 1, while *R*_0_ may be maximized at lower values, thus dampening selection towards high virulence. The resulting increase in virulence then leads to higher host investment into resistance.

### Inducible defense creates an epidemiological feedback that determines host resistance

The force of infection in our model is mainly driven by the parasite dispersal rate *m*. When parasite dispersal is close to the population viability boundaries (respectively low or high enough, see fig.(2) in Appendix A), the force of infection is weak. While parasite levels of investment remain high across a wide range of intermediate dispersal values, host investment is more variable (fig 4). As a general result, we find that the host’s investment in resistance and the force of infection follow an inverse pattern.

**Figure 3.**
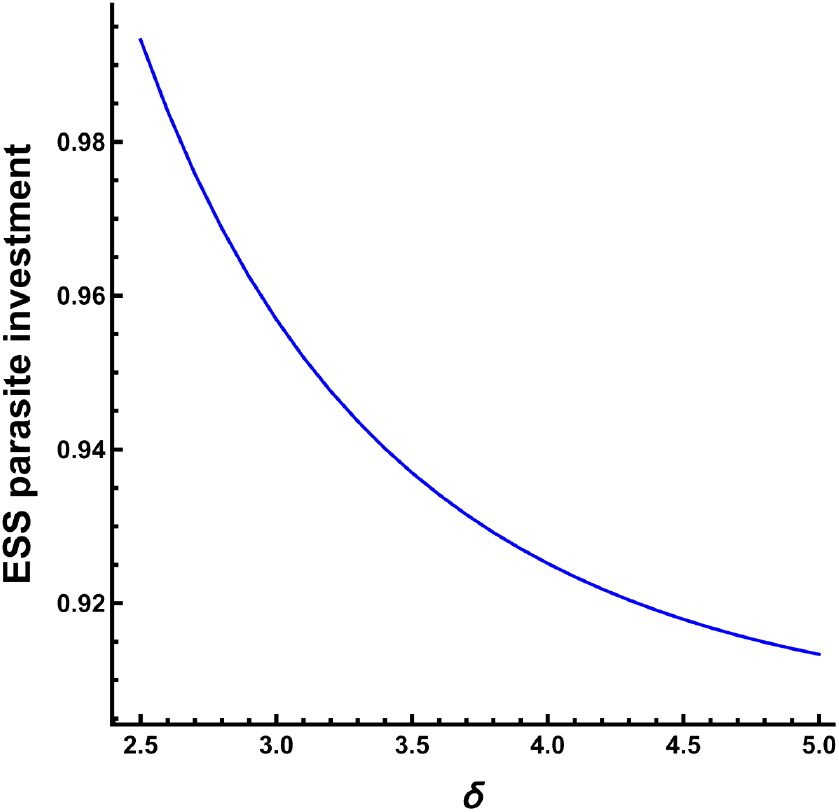
CSS Parasite virulence as a function of trade-off shape between virulence and replication, with *θ* left as a variable and *σ* set to 0.5. *δ* defines the shape of the trade-off function (concave when *δ <* 1, linear when *δ* = 1, convex when *δ >* 1. Other parameters were fixed to *b*_0_ = 0.8, *µ* = 0.5, *α* = 1, *r*_0_ = 10, *r*_1_ = 0.04, *ρ* = 0.9, *γ*_0_ = 1, *d*_0_ = 0.05, *τ* = 0.5, *ϵ* = 3, *κ* = 1, *d* = 0.5

**Figure 4.**
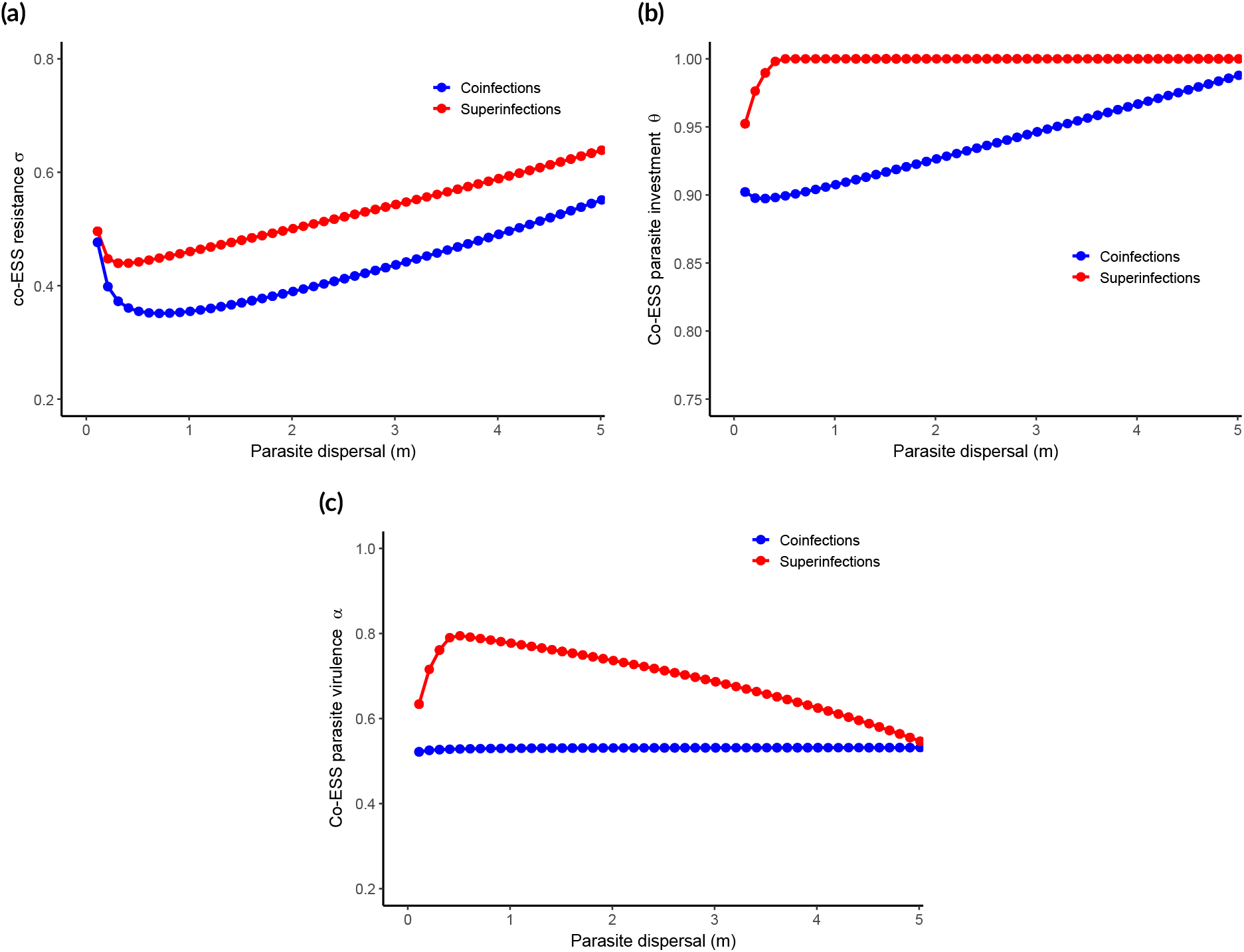
Coevolutionary singular strategies as a function of parasite dispersal rate *m* for the host resistance *σ*, b) the parasite investment *θ* and c) the parasite virulence *α*. Red dots refers to results derived from the superinfection model, while blue refers to results derived from the coinfection model. Fixed parameters were set to :*ϵ* = 5, *b*_0_ = 0.8, *µ* = 0.5, *α*_0_ = 0.5, *r*_0_ = 10,, *r*_1_ = 0.04, *ρ* = 0.9, *γ*_0_ = 1, *d*_0_ = 0.05, *τ* = 1, *δ* = 5, *κ* = 1, *η* = 1.

When parasite dispersal is very low, host resistance is relatively high, although it decreases at first as dispersal increases. In this regime, the disease is rare but infections are severe, so resistance is favored because its costs are paid by few individuals. Low dispersal also makes multiple infections rare, so coinfection and superinfection give similar results and approach the single-infection case (Fig. 4a).

When dispersal approaches its upper viability limit, the force of infection is low because dispersal is costly. Few hosts are infected, and parasite loads remain small. Host resistance is then favored, which increases recovery and reduces mortality risk (Fig. 4c). In response, the parasite invests more in replication.

When dispersal is intermediate, infection is common. In the superinfection case, parasite investment quickly reaches its maximum, which increases parasite load and shortens infections. At the same time, host investment keeps increasing, which lowers virulence and pushes the parasite toward the *R*_0_ optimization principle by extending the infectious period (Fig. 4b; Fig. 4c; Fig. 5c). Under coinfection, both host and parasite investment increase with dispersal, but the high cost of dispersal offsets these effects, so virulence rises only slightly (Fig. 4c).

**Figure 5.**
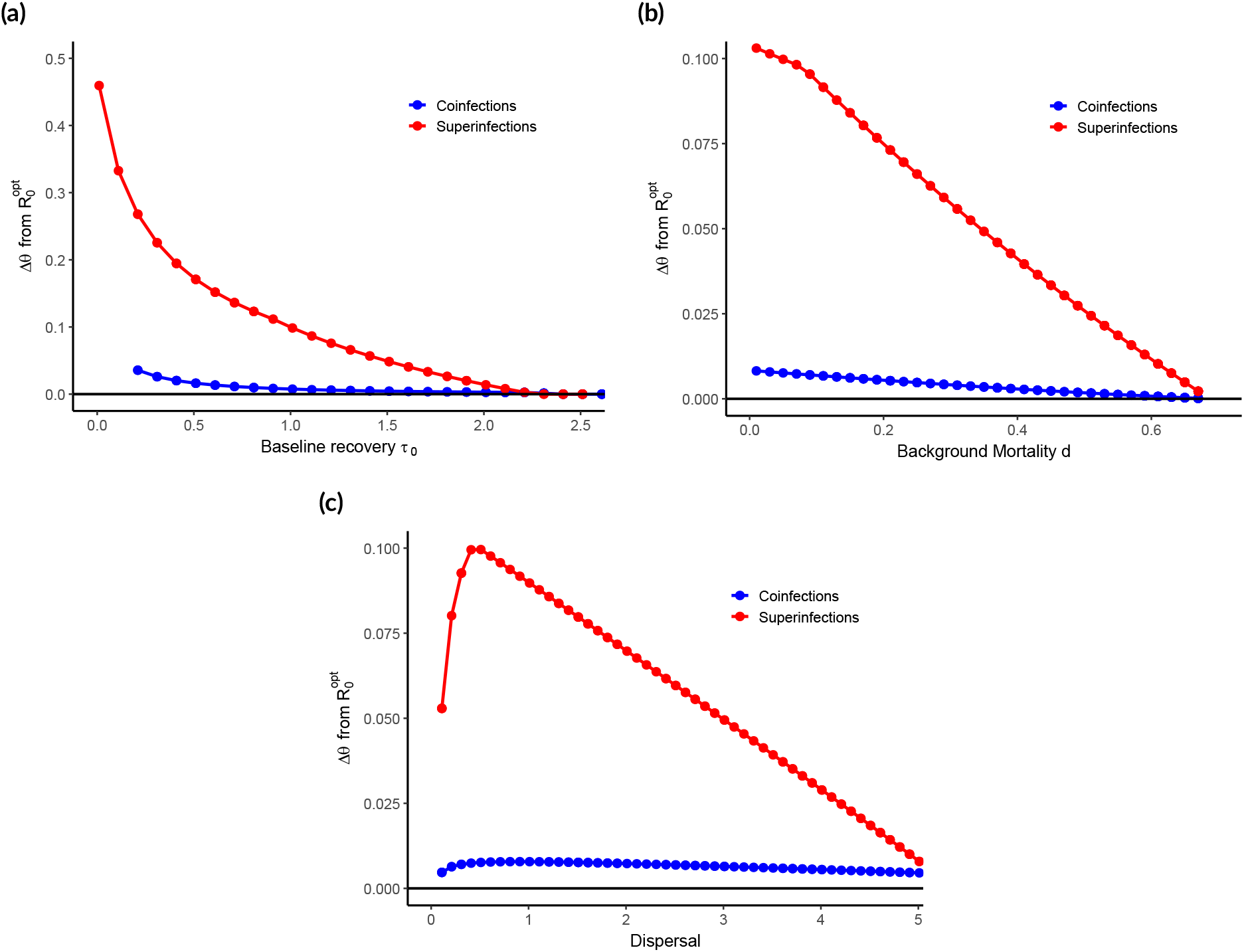
Deviation between the coevolved *R*_0_ and the *R*_0_ predicted by an optimization principle, evaluated at the host co-singular strategy, in the coinfection model (blue curve) and superinfection model (red curve). Constant parameters were set to *b*_0_ = 1, *µ* = 0.5, *r*_0_ = 10, *r*_1_ = 0.04, *ρ* = 0.9, *d*_0_ = 0.05, *τ* = 1, *d*_0_ = 0.05, *ϵ* = 5, *α* = 1, *γ*_0_ = 1, *δ* = 5, *κ* = 1, *η* = 1. (a) Deviation is plotted according to the baseline recovery rate *τ*_0_. (b) Deviation is plotted according to the host background mortality *d*.(c) Deviation is plotted according to the parasite dispersal rate *m*

These results are a consequence of our assumption made on the distribution of resistance costs. We assumed that only infected individuals pay the costs of resistance, and considered that the cost acts on host fecundity. Doing so, the force of infection creates a negative demographic feedback on host population through the infection prevalence. Selection on resistance therefore generally leads the host towards less investment when the disease is widespread and sufficiently lethal.

Our assumption that resistance costs are induced also affects how host and parasite investments respond to changes in recovery rate. Figures 6a and 6b show that higher recovery selects for both greater parasite replication and greater resistance, with a stronger effect in the super-infection case. This is consistent with previous work showing that increased recovery can favor higher parasite exploitation van Baalen, 1998. In our model, recovery benefits the host because recovered individuals return to the susceptible class and no longer pay resistance costs. A higher baseline recovery also shortens the infectious period, which reduces the time during which those costs are paid. As a result, resistance is favored because it further shortens infection and reduces virulence (Fig. 6c). This is consistent with previous theory suggesting that inducible defenses are favored when infection is rare or when their costs are low Cressler et al., 2015.

**Figure 6.**
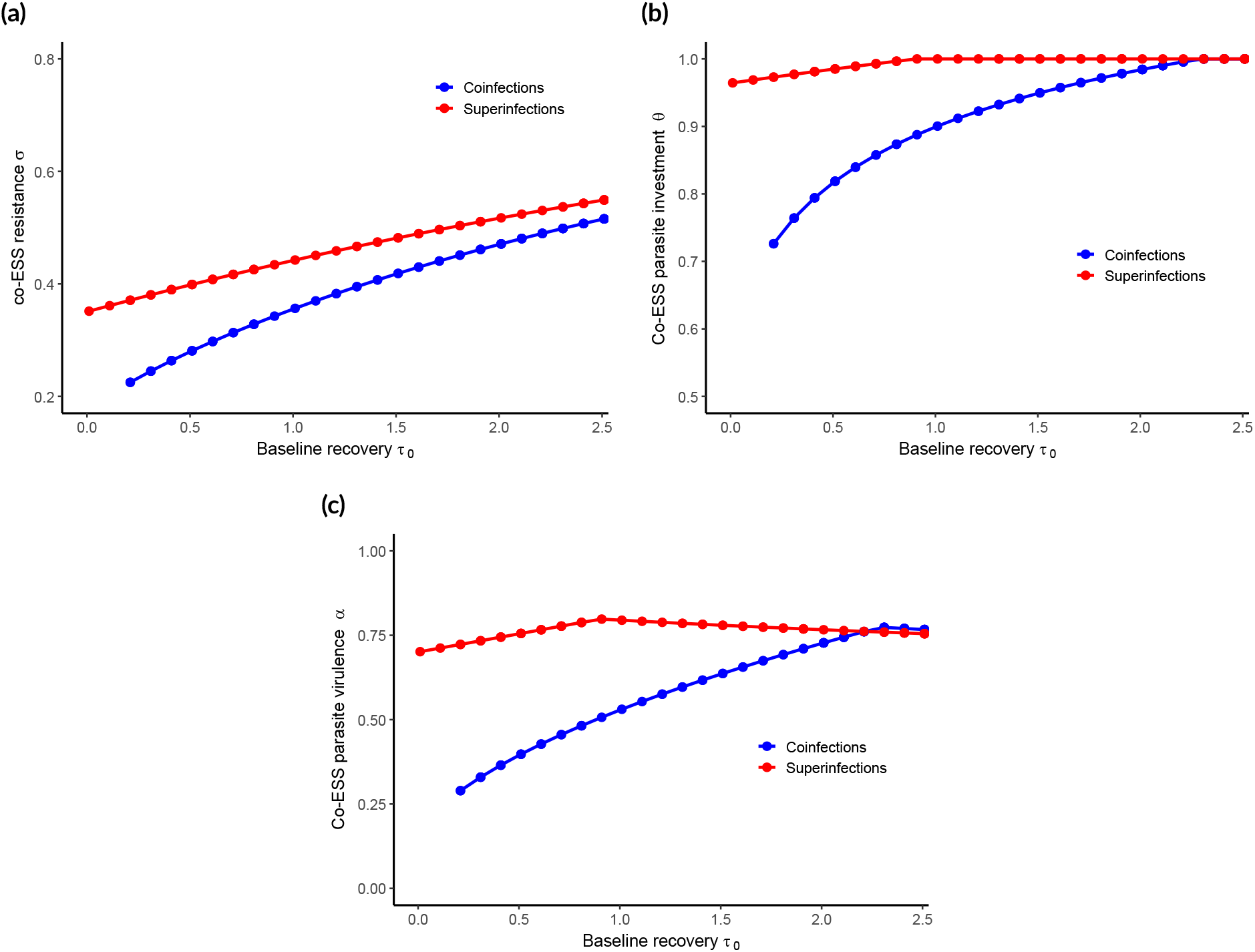
Coevolutionary singular points as a function of host baseline recovery rate for a) the host resistance *σ*, b) the parasite investment *θ* and c) the parasite virulence *α*. Red lines refers to results derived from the superinfection model, while blue lines refers to results derived from the coinfection model. Fixed parameters were set to :*ϵ* = 5, *b*_0_ = 0.8, *µ* = 0.5, *α* = 1, *r*_0_ = 10, *r*_1_ = 0.04, *ρ* = 0.9, *γ*_0_ = 1, *d*_0_ = 0.05, *δ* = 5, *κ* = 1, *d* = 0.5, *η* = 1.

### Deviation from *R*_0_ optimization emerges from a competition-colonization trade-off

Host investment in resistance according to the host background mortality (fig. 7a) shows a quite intuitive pattern. As the host lifespan diminishes, so does the investment in resistance. An intuitive explanation is that the host fecundity is favored over resistance to infectious agents in short-lived species (Miller et al., 2007). However, the parasite investment and virulence seems, at first glance, more counter-intuitive (figs. 7b,7c). While virulence seems favored when the host lifespan is short in the case of coinfections, superinfections show the opposite tendency. Such a result can however be understood in terms of optimization principles.

**Figure 7.**
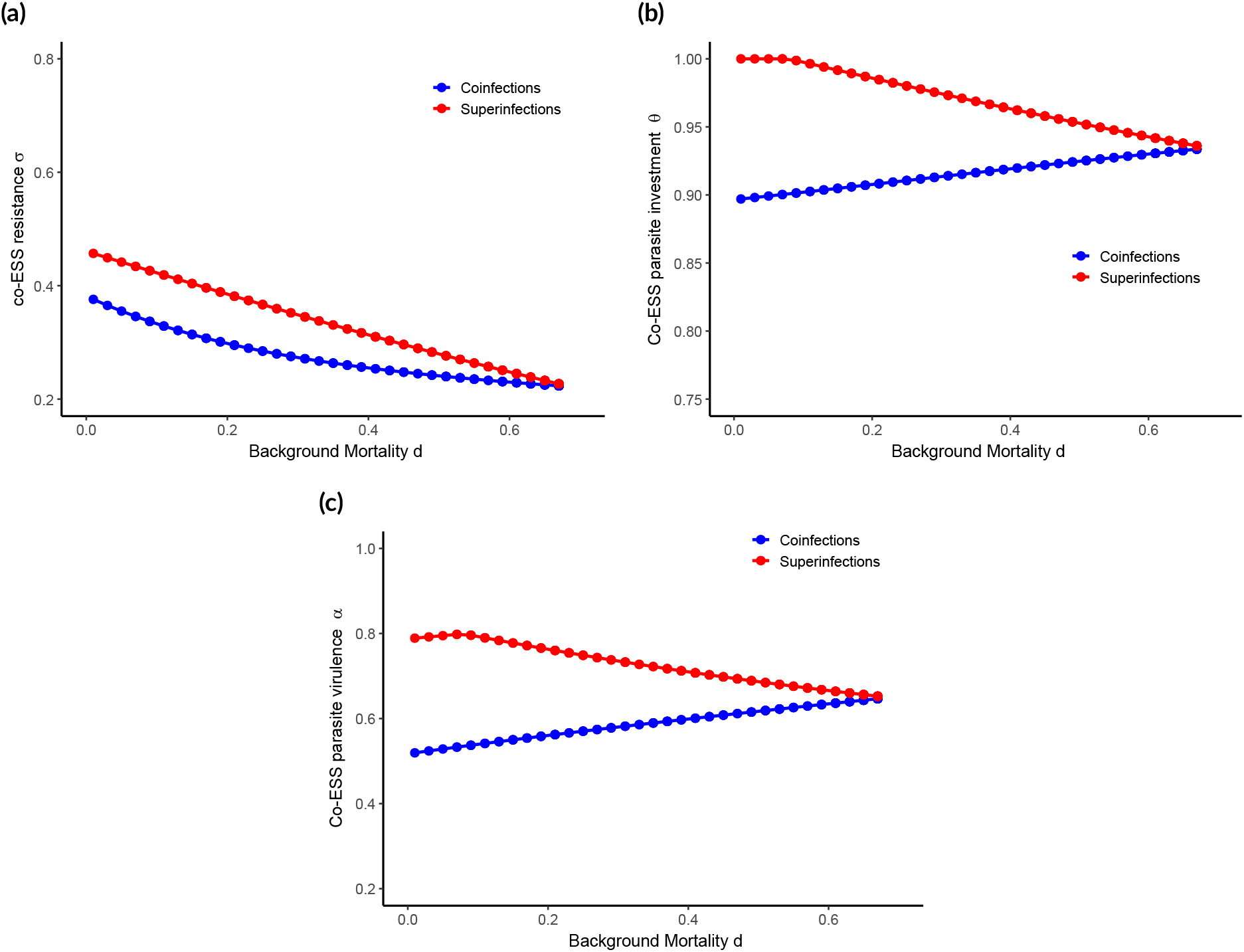
Coevolutionary singular points as a function of host background mortality for a) the host resistance *σ*, b) the parasite investment *θ* and c) the parasite virulence *α*.. Red lines refers to results derived from the superinfection model, while blue lines refers to results derived from the coinfection model. Fixed parameters were set to : *ϵ* = 5, *b*_0_ = 0.8, *µ* = 0.5, *α* = 1, *r*_0_ = 10, *r*_1_ = 0.04, *ρ* = 0.9, *γ*_0_ = 1, *τ*_0_ = 1, *δ* = 5, *κ* = 1, *d* = 0.5, *η* = 1.

Simple models of parasite evolution state that virulence should be selected towards values that maximizes the basic reproductive ratio *R*_0_ (Lion and Metz, 2018). The addition of complexity (*i*.*e*. multiple infections and host evolution) is expected to cause deviation from the *R*_0_ optimization principle, as it increases the dimensionality of the environmental feedback loop. We have shown that when only the parasite evolves, virulence should evolve towards higher levels in multiple infection than expected under the single infection scenario, as a result of strain competition (eqn. 17). Host evolution does not change this expectation (fig 5). However, we also find that the *R*_0_ optimization principle holds in our model when the host or the parasite are driven towards their viability limit. This can be explained by the reduction of the environmental feedback loop, which can be seen as a consequence of a scenario in which trait evolution is driven by ecological persistence. Near to the viability boundary, the occurence of multiple infection tends to be drastically reduced because when infections are scarce, double infections are scarcer (*i*.*e*. 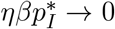 in eqn.15). Thus the environmental feedback loop acting on the parasite should tend to be only determined by the infection of susceptible individuals.

On the contrary, when the model parameters are set far from the parasite viability boundary (intermediate dispersal (m), low baseline recovery *τ*_0_ or low background mortality *d*), we have shown that dominance competition has a larger contribution to parasite fitness (eqn. 17, fig. 5). Competitive ability in our model is acquired at the cost of an increased virulence, which decreases the within-host persistence time. Thus, parasite experience two contradictory selective pressures. The first is expected to maximize the competitive ability, and therefore the parasite replication rate *r*_0_. The maximization of *r*_0_ thus correspond to the maximization of the within-host equilibrium load *X*^∗^, and consequently to maximization of the within-host competitive ability. The second selective pressure is the one that is expected to maximize the basic reproductive number *R*_0_ which, from a metapopulation perspective, can be understood here as the rate of successful colonization of new patches by the parasite.

As soon as the value of *θ* that optimizes *r*_0_ differs from the one that optimizes *R*_0_ (or equivalently, when 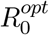 does not correspond to *θ* = 1), conflicting selection induces the emergence of a competition - colonization trade-off (Messinger and Ostling, 2009). Not surprisingly, this trade-off is more intense in the superinfection model than in the coinfection model. As superinfections assumes no preemptive effects in strain competition, this enables a stricter competitive hierarchy in parasite, because competitive exclusion always occurs in infected hosts.

Allowing for some preemption through coinfections only leads to a slight deviation from the *R*_0_ optimization. The addition of preemption implies the local coexistence of strains, because exclusion may not have occurred at the end of infection. The weak deviation from models that obey to this optimization principle is thus explained by two features that stem from our coinfection framework : (i) the assumption of weak competitive differences, that mechanically dampen the intensity of dominance competition and (ii) the induction of a kin selection effect which always drives selection on virulence in the opposite direction to competition.

## Discussion

Our work aims to bridge the gap existing between the ecology of subdivided populations, and theoretical epidemiology. Following an approach initiated by Restif and Koella (2003), we built an epidemiological model under the assumption that virulence, transmission and recovery are primarily characteristics of a disease, which results from the interaction between the individual traits of a host and its parasite. Using the parasite load as a proxy affecting these three main epidemic features, we developed a modelling framework rooted in metapopulation ecology to assess the host and parasite coevolution, in the context of multiple infections, and under different regimes of competition.

Our broader result is that the competition regime strongly affects the evolution of parasite virulence, and determines the host response to the selective pressures induced by the ecological conditions. When strain competition acts mainly through dominance, virulence evolves towards higher values, and the host generally responds by an increased investment in resistance. However, the addition of preemptive effects in strain competition reduces the overall harm caused by the parasite. Such results are the consequence of two major properties arising from our modelling framework.

First, a competition-colonization (hereafter CC) trade-off that emerges from the conflictual selective pressures between the colonization of new susceptibles and the within-host competitiveness (Mideo, 2009). The first is expected to maximize the epidemiological *R*_0_, while the second is maximized with *r*_0_*θ*, at the cost of a reduction in local persistence time (Messinger and Ostling, 2009). Such a trade-off is representative of spatially structured populations (Tilman, 1994), and emerges in our model from limited dispersal between discrete host patches.

Second, a kin selection effect that is induced by the local coexistence of strains, and acts in an opposite direction to competition. Relatedness in coinfecting parasites have been shown to lead to various outcomes according to the nature of exploitation behaviour of individuals (Brown et al., 2002, Buckling and Brockhurst, 2008). In our model, virulence is a consequence of the collective exploitation of the host patch by the parasite population(s), and depends on both individual traits and the total parasite load. Our work provides results that are consistent with the idea that when exploitation is limited by the collective action of individuals, relatedness should favor prudent strategies (although it may also depend on the determinism and nature of the considered within-host interactions, *e*.*g*. Alizon and Lion, 2011).

A corollary result is that both the CC trade-off and the kin selection effect vary depending on the ecological context. In particular, we have shown that both the parasite transmission success and host demography determine the existence of the CC trade-off (fig. 5). Successful dispersal of parasite propagules can be seen as the degree of coupling between host patches, and reflects the contact intensity in the host population. Such coupling determines the dimensions of the eco-evolutionary feedback loop acting on the parasite. When the infection is rare, the (eco-to-evo) feedback drives selection towards maximized persistence because transmission occurs mostly from infected to suceptibles. Conversely, widespread parasites cause infected individuals to act as a supplementary ecological niche. This causes deviation from the simple optimization of persistence through the epidemiological *R*_0_, which is a classic result obtained from the study of multiple infections (Mosquera and Adler, 1998, Alizon and van Baalen, 2008).

We have shown that when coevolution leads to reduced levels of parasite exploitation, the host generally invests less in resistance. This is not surprising, since the damages induced by the parasite are then also reduced. More importantly, this result helps reinterpret what is expected from the single-infection model when epidemiological traits are under shared control. In the absence of multiple infections, host investment in resistance is expected to peak at intermediate parasite virulence Restif and Koella, 2003. We did not retrieve this behavior, and we argue that this is due to the epidemiological and demographic feedbacks generated by the force of infection 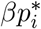, which themselves depend on how resistance costs are distributed. Because Restif and Koella (2003) assumed a constitutive cost of resistance, infection does not change host fecundity in their model, so the population-level cost of resistance remains unchanged when the force of infection varies. In our model, by contrast, the force of infection determines prevalence and therefore the population-level magnitude of resistance costs, since only infected individuals suffer reduced fecundity. As a result, resistance can become counter-selected when parasites become ubiquitous in the population. Such a finding remains consistent with the general idea provided by most (if not all) theoretical models: resistance is counter-selected as soon as its costs exceed its benefits. That said, whether the cost exceeds the benefits depends on: i) the action mode of the trait (*i*.*e*. avoidance, clearance), ii) the action mode of the cost (*i*.*e*. induced, constitutive, or both) and iii) the eco-epidemiological feedback generated by the two previous ingredients Boots and Haraguchi, 1999; Best et al., 2010; Donnelly et al., 2015; Singh and Best, 2023. In our model, resisting when surrounded by “enemies” would increase the demographic depression already caused by the parasite through the diminished reproduction of infected hosts that are, in any case, unable to clear their parasite. This phenomenon, although rarely highlighted in theoretical literature, was mathematically detailed and experimentally observed by Walsman et al. (2023).

On the other hand, the infectious period (and its consequences on the host demography) is also an important driver of the eco-evolutionary feedback loop. In addition, its length shapes the balance between preemptive and dominance competiiton.

When a reduced infectious period results in a positive demographic feeback on the host population (*e*.*g*. by an increase in recovery), only the parasite is pushed towards its viability limit. Then, we predict that selection should favor higher virulence and resistance (figs. 6) than observed with varying background mortality (figs. 7). This results from both the shared control of epidemic features, and our assumption on the induced nature of resistance. High resistance allows the host to dampen the deleterious effects of the parasite by reducing both transmission and mortality risks, which is a consequence of the shared control. At the same time, increased resistance enhances recovery, which has twofold benefits, because a recovered host suffers neither from parasite effects, nor from the costs of resistance. The cost of resistance is thus sufficiently cheap to favor investment (Cressler et al., 2015). Conversely, when the demographic feedback of a reduced infectious period is negative (*i*.*e*. by an increase in host mortality), both the host and the parasite are driven towards their viability limit. Selection here leads the traits evolution towards persistence, which correspond to low (but non-zero) resistance (fig. 7a).

The expression of parasite’s fitness we derived (eq. 15) is conceptually interesting because it enables a straightforward link between several theoretical approaches. By considering a well-mixed host population as a set of discrete patches coupled through parasite dispersal, we showed that the measure of the lifetime population success *R*_*m*_ (Metz and Gyllenberg, 2001) is equivalent to the reproductive numbers traditionally used in theoretical epidemiology in the absence of multiple infections (Lion and Metz, 2018, Almocera et al., 2018). Our nested approach also emphasizes the now recognized links between fitness in subdivided populations and inclusive fitness theory (Ajar, 2003), as well as the vision of the altruistic nature of virulence-related traits (Griffin et al., 2004). Moreover, it provides a natural framework for multiscale modelling, accounting for a variety of local interactions and the joint evolution of multiple epidemic quantities, which are often treated independently. In addition, the links between the different interaction scales, as well as the evolving individual traits, are defined through quantities that are commonly measured in life sciences (*e*.*g*. parasite load, division rate or clearance rate following phagocytosis by macrophages). We believe our methodological framework could thus help empiricists and theoreticians to design system-specific models and experiments to refine our mechanistic understanding of host-parasite interactions.

Another strength of our model is that the emergence of multi-scale selection as well as demographic feedbacks are not affected by the assumptions made on the cost profile between host and parasite traits (*i*.*e*. our eqn.(17) is blind to the trade-off shapes). However, some other results are expected to be sensitive to the assumptions made during our analysis. It is now well-recognized that the choice of the ‘take-over’ function in multiple infection models can have significant effects on the qualitative outcome of the coevolutionary dynamics, especially relative to the emergence of stable polymorphism (Mosquera and Adler, 1998; Boldin and Diekmann, 2008). By choosing to set *η* = 1 in our numerical analysis, we assumed that any secondary invasion of a host was by definition successful. As a result, an undercompetitive mutant remains capable of producing propagules for a non-zero period of time. Moreover, the simple fact that having defined *η* as a fixed parameter is a limitation in itself, because such a deterministic behaviour is quite unlikely to occur in natural systems. A natural expansion of our model would account for the early stochastic dynamics of invasion (*e*.*g*. by defining the probability of multiple infections as a function of competitive traits (see Kada and Lion, 2015; Boldin and Diekmann, 2008 for examples).

Another limitation of our model relies on the assumptions made on the interaction between host defense and parasite growth. We chose a simple model of parasite growth that accounts for parasite clearance and derived several mechanisms of epidemic mitigation and infection clearance at the host population scale. No possibility of tolerance mechanisms were accounted for. This may have important implications for host and parasite coevolution (Carval and Ferriere, 2010, Best et al., 2010) especially in the case of multiple infections. Tolerance is defined in theoretical works as having a positive effect on parasite fitness (Carval and Ferriere, 2010), and has been shown to be traded off with resistance in several systems (Råberg et al., 2007; Salgado-Luarte et al., 2023). Recent studies have shown this particular trade-off may strongly affect modes of selection (directional or fluctuating) and alter the possibility of stable coevolutionary coexistence (Singh, 2023). We expect such considerations to greatly affect our results.

The within-host dynamics chosen in our study does not capture the whole diversity of parasite interactions occurring in natural systems, such as production of public goods, or spiteful behaviours (Leinweber et al., 2018; Niehus et al., 2017; Bucci et al., 2011; Bashey et al., 2012). Strain competition in our model acts through (i) exploitation competition for the exact same host resources and (ii) apparent competition against the host’s immune system, which can be seen as acting like a generalist predator. However, parasite evolution in our work only accounts for differences in the intensity of exploitation competition with no possibility to avoid apparent competition. In natural systems, this is achieved through adaptation towards shifts in antigenic determinants that allows new mutant to escape from host immune response (Fryer et al., 2010). It would be interesting to take into account other patterns of within-host parasite interactions in future work.

Following previous works on multiple infections, we use a quasi-equilibrium assumption to describe the variation in within-host parasite density during competition. Events of co-transmission, or time heterogeneity in the sequence of invasions by multiple parasites for example, would certainly lead to different competitive outcomes, subsequently affecting the coevolutionary dynamics and associated feedbacks. Empirical evidence for such alterations have been reviewed in (Dutt et al., 2022), and are also suggested by contradictory results obtained by clinical studies (Sullivan et al., 2015, Fry et al., 2019). One of the crucial challenges for future theoretical studies assessing how parasite competition would alter coevolutionary dynamics will rely on our ability to describe more complex interactions, and address how the ‘age at secondary invasion’ affects the outcome of parasite competition.

## Acknowledgements

We thank Jan Engelstaedter and two anonymous reviewers for very useful comments on a previous version of this manuscript.

## Funding

This study is part of the FEEDME project coordinated by O. Kaltz, and has been funded by a grant from the Agence Nationale de la Recherche (NO : ANR-20-CE02-0023) to F. Massol and S. Lion.

## Conflict of interest disclosure

The authors declare that they comply with the PCI rule of having no financial conflicts of interest in relation to the content of the article.

## Data, script, code, and supplementary information availability

Datas, script, codes and supplementary information are available online (https://doi.org/10.5281/zenodo.12755175).

## Notes

https://doi.org/10.5281/zenodo.12755175

### Competing Interest Statement

The authors have declared no competing interest.

### Summary of Updates

Minor changes related to document formatting and typo correction have been made. All the main text figures have been redrawn in a more esthetic way, and new ones are provided, according to reviewers's specific comments.

## References

Acevedo MA, Dillemuth FP, Flick AJ, Faldyn MJ, Elderd BD (2019). Virulence-driven trade-offs in disease transmission: A meta-analysis. Evolution 73, 636–647. 10.1111/evo.13692.

Ajar É (2003). Analysis of disruptive selection in subdivided populations. BMC Evolutionary Biology 3, 1–12. 10.1186/1471-2148-3-22.

Alizon S, De Roode JC, Michalakis Y (2013). Multiple infections and the evolution of virulence. Ecology letters 16, 556–567. 10.1111/ele.12076.

Alizon S, Hurford A, Mideo N, van Baalen M (2009). Virulence evolution and the trade-off hypothesis: history, current state of affairs and the future. Journal of evolutionary biology 22, 245–259. 10.1111/j.1420-9101.2008.01658.x.

Alizon S, Lion S (2011). Within-host parasite cooperation and the evolution of virulence. Proceedings of the Royal Society B: Biological Sciences 278, 3738–3747. 10.1098/rspb.2011.0471.

Alizon S, van Baalen M (2005). Emergence of a convex trade-off between transmission and virulence. The American Naturalist 165, E155–E167. 10.1086/430053.

Alizon S, van Baalen M (2008). Multiple infections, immune dynamics, and the evolution of virulence. The American Naturalist 172, E150–E168. 10.1086/590958.

Almocera AES, Nguyen VK, Hernandez-Vargas EA (2018). Multiscale model within-host and between-host for viral infectious diseases. Journal of Mathematical Biology 77, 1035–1057. 10.1007/s00285-018-1241-y.

Bashey F, Young S, Hawlena H, Lively C (2012). Spiteful interactions between sympatric natural isolates of Xenorhabdus bovienii benefit kin and reduce virulence. Journal of evolutionary biology 25, 431–437. 10.1111/j.1420-9101.2011.02441.x.

Best A, Long G, White A, Boots M (2012). The implications of immunopathology for parasite evolution. Proceedings of the Royal Society B: Biological Sciences 279, 3234–3240.

Best A, White A, Boots M (2009). The implications of coevolutionary dynamics to host-parasite interactions. The American Naturalist 173, 779–791. 10.1086/598494.

Best A, White A, Boots M (2010). Resistance is futile but tolerance can explain why parasites do not always castrate their hosts. Evolution 64, 348–357. 10.1111/j.1558-5646.2009.00819.x.

Boldin B, Diekmann O (2008). Superinfections can induce evolutionarily stable coexistence of pathogens. Journal of mathematical biology 56, 635–672. 10.1007/s00285-007-0135-1.

Boots M, Haraguchi Y (1999). The evolution of costly resistance in host-parasite systems. The american naturalist 153, 359–370. 10.1086/303181.

Brown SP, Hochberg ME, Grenfell BT (2002). Does multiple infection select for raised virulence? Trends in microbiology 10, 401–405. 10.1016/s0966-842x(02)02413-7.

Bucci V, Nadell CD, Xavier JB (2011). The evolution of bacteriocin production in bacterial biofilms. The American Naturalist 178, E162–E173. 10.1086/662668.

Buckling A, Brockhurst M (2008). Kin selection and the evolution of virulence. Heredity 100, 484–488. 10.1038/sj.hdy.6801093.

Calcagno V, Mouquet N, Jarne P, David P (2006). Coexistence in a metacommunity: the competition–colonization trade-off is not dead. Ecology letters 9, 897–907. 10.1111/j.1461-0248.2006.00930.x.

Carval D, Ferriere R (2010). A unified model for the coevolution of resistance, tolerance, and virulence. Evolution 64, 2988–3009. 10.1111/j.1558-5646.2010.01035.x.

Cressler CE, Graham AL, Day T (2015). Evolution of hosts paying manifold costs of defence. Proceedings of the Royal Society B: Biological Sciences 282, 20150065. 10.1098/rspb.2015.0065.

Cressler CE, McLeod DV, Rozins C, Van Den Hoogen J, Day T (2016). The adaptive evolution of virulence: a review of theoretical predictions and empirical tests. Parasitology 143, 915–930. 10.1017/S003118201500092X.

Day T, Alizon S, Mideo N (2011). Bridging scales in the evolution of infectious disease life histories: theory. Evolution 65, 3448–3461. 10.1111/j.1558-5646.2011.01394.x.

De Roode JC, Helinski ME, Anwar MA, Read AF (2005). Dynamics of multiple infection and within-host competition in genetically diverse malaria infections. The American Naturalist 166, 531–542. 10.1086/491659.

Diekmann O, Heesterbeek JAP, Metz JA (1990). On the definition and the computation of the basic reproduction ratio R 0 in models for infectious diseases in heterogeneous populations. Journal of mathematical biology 28, 365–382. 10.1007/BF00178324.

Donnelly R, White A, Boots M (2015). The epidemiological feedbacks critical to the evolution of host immunity. Journal of Evolutionary Biology 28, 2042–2053. 10.1111/jeb.12719.

Dutt A, Andrivon D, Le May C (2022). Multi-infections, competitive interactions, and pathogen co-existence. Plant Pathology 71, 5–22. 10.1111/ppa.13469.

Eshel I (1983). Evolutionary and continuous stability. Journal of theoretical Biology 103, 99–111.

Frank SA (1996). Models of parasite virulence. The Quarterly review of biology 71, 37–78. 10.1086/419267.

Frank SA (2012). Natural selection. III. Selection versus transmission and the levels of selection. Journal of evolutionary biology 25, 227–243. 10.1111/j.1420-9101.2011.02431.x.

Frickel J, Sieber M, Becks L (2016). Eco-evolutionary dynamics in a coevolving host–virus system. Ecology letters 19, 450–459. 10.1111/ele.12580.

Fry SHL, Barnabas SL, Cotton MF (2019). Tuberculosis and HIV—An Update on the “cursed duet” in children. Frontiers in pediatrics 7, 159. 10.3389/fped.2019.00159.

Fryer HR, Frater J, Duda A, Roberts MG, Investigators ST, Phillips RE, McLean AR (2010). Modelling the evolution and spread of HIV immune escape mutants. PLoS pathogens 6, e1001196. 10.1371/journal.ppat.1001196.

Geritz SA, Kisdi E, Mesze NA G, Metz JA (1998). Evolutionarily singular strategies and the adaptive growth and branching of the evolutionary tree. Evolutionary ecology 12, 35–57. 10.1023/A:1006554906681.

Griffin AS, West SA, Buckling A (2004). Cooperation and competition in pathogenic bacteria. Nature 430, 1024–1027. 10.1038/nature02744.

Hastings A (1980). Disturbance, coexistence, history, and competition for space. Theoretical population biology 18, 363–373. 10.1016/0040-5809(80)90059-3.

Hurford A, Cownden D, Day T (2010). Next-generation tools for evolutionary invasion analyses. Journal of the Royal Society Interface 7, 561–571. 10.1098/rsif.2009.0448.

Jansen VA, Vitalis R (2007). THE EVOLUTION OF DISPERSAL IN A LEVINS’TYPE METAPOPULATION MODEL. Evolution 61, 2386–2397. 10.1111/j.1558-5646.2007.00201.x.

Kada S, Lion S (2015). Superinfection and the coevolution of parasite virulence and host recovery. Journal of evolutionary Biology 28, 2285–2299. 10.1111/jeb.12753.

Kermack WO, McKendrick AG (1927). A contribution to the mathematical theory of epidemics. Proceedings of the royal society of london. Series A, Containing papers of a mathematical and physical character 115, 700–721. 10.1098/rspa.1927.0118.

Lefèvre T, Sanchez M, Ponton F, Hughes D, Thomas F (2007). Virulence and resistance in malaria: who drives the outcome of the infection? Trends in parasitology 23, 299–302. 10.1016/j.pt.2007.04.012.

Leinweber A, Weigert M, Kümmerli R (2018). The bacterium Pseudomonas aeruginosa senses and gradually responds to interspecific competition for iron. Evolution 72, 1515–1528. 10.1111/evo.13491.

Levins R (1969). Some demographic and genetic consequences of environmental heterogeneity for biological control. Bulletin of the ESA 15, 237–240. 10.1093/besa/15.3.237.

Lion S (2013). Multiple infections, kin selection and the evolutionary epidemiology of parasite traits. Journal of evolutionary biology 26, 2107–2122. 10.1111/jeb.12207.

Lion S, Metz JA (2018). Beyond R0 maximisation: on pathogen evolution and environmental dimensions. Trends in ecology & evolution 33, 458–473. 10.1016/j.tree.2018.02.004.

Massol F, Calcagno V, Massol J (2009). The metapopulation fitness criterion: proof and perspectives. Theoretical population biology 75, 183–200. 10.1016/j.tpb.2009.02.005.

May RM, Anderson RM (1979). Population biology of infectious diseases: Part II. Nature 280, 455–461. 10.1038/280455a0.

May RM, Nowak MA (1995). Coinfection and the evolution of parasite virulence. Proceedings of the Royal Society of London. Series B: Biological Sciences 261, 209–215. 10.1098/rspb.1995.0138.

Messinger SM, Ostling A (2009). The consequences of spatial structure for the evolution of pathogen transmission rate and virulence. The American Naturalist 174, 441–454. 10.1086/605375.

Metz J, Gyllenberg M (2001). How should we define fitness in structured metapopulation models? Including an application to the calculation of evolutionarily stable dispersal strategies. Proceedings of the Royal Society of London. Series B: Biological Sciences 268, 499–508. 10.1098/rspb.2000.1373.

Mideo N (2009). Parasite adaptations to within-host competition. Trends in parasitology 25, 261–268. 10.1016/j.pt.2009.03.001.

Mideo N, Alizon S, Day T (2008). Linking within-and between-host dynamics in the evolutionary epidemiology of infectious diseases. Trends in ecology & evolution 23, 511–517. 10.1016/j.tree.2008.05.009.

Miller MR, White A, Boots M (2007). Host life span and the evolution of resistance characteristics. Evolution 61, 2–14. 10.1111/j.1558-5646.2007.00001.x.

Mosquera J, Adler FR (1998). Evolution of virulence: a unified framework for coinfection and super-infection. Journal of Theoretical Biology 195, 293–313. 10.1006/jtbi.1998.0793.

Niehus R, Picot A, Oliveira NM, Mitri S, Foster KR (2017). The evolution of siderophore production as a competitive trait. Evolution 71, 1443–1455. 10.1111/evo.13230.

Nowak MA, May RM (1994). Superinfection and the evolution of parasite virulence. Proceedings of the Royal Society of London. Series B: Biological Sciences 255, 81–89. 10.1098/rspb.1994.0012.

Otto SP, Day T (2007). A biologist’s guide to mathematical modeling in ecology and evolution. Princeton University Press.

Pillai P, Gonzalez A, Loreau M (2012). Evolution of dispersal in a predator-prey metacommunity. The American Naturalist 179, 204–216. 10.1086/663674.

Råberg L, Sim D, Read AF (2007). Disentangling genetic variation for resistance and tolerance to infectious diseases in animals. Science 318, 812–814. 10.1126/science.1148526.

Restif O, Koella JC (2003). Shared control of epidemiological traits in a coevolutionary model of host-parasite interactions. The American Naturalist 161, 827–836. 10.1086/375171.

Rousset, Billiard (2000). A theoretical basis for measures of kin selection in subdivided populations: finite populations and localized dispersal. Journal of Evolutionary Biology 13, 814–825. 10.1046/j.1420-9101.2000.00219.x.

Salgado-Luarte C, González-Teuber M, Madriaza K, Gianoli E (2023). Trade-off between plant resistance and tolerance to herbivory: Mechanical defenses outweigh chemical defenses. 10.1002/ecy.3860.

Singh P (2023). Trade-offs between host defense mechanisms: impacts on evolutionary and coevolutionary dynamics of host-parasite interactions. PhD thesis. University of Sheffield.

Singh P, Best A (2023). A Sterility–Mortality Tolerance Trade-Off Leads to Within-Population Variation in Host Tolerance. Bulletin of Mathematical Biology 85, 16. 10.1007/s11538-023-01119-6.

Sofonea MT, Alizon S, Michalakis Y (2015). From within-host interactions to epidemiological competition: a general model for multiple infections. Philosophical Transactions of the Royal Society B: Biological Sciences 370, 20140303.

Sullivan ZA, Wong EB, Ndung’u T, Kasprowicz VO, Bishai WR (2015). Latent and active tuberculosis infection increase immune activation in individuals co-infected with HIV. EBioMedicine 2, 334–340. 10.1016/j.ebiom.2015.03.005.

Susi H, Vale PF, Laine AL (2015). Host genotype and coinfection modify the relationship of within and between host transmission. The American Naturalist 186, 252–263. 10.1086/682069.

Tilman D (1994). Competition and biodiversity in spatially structured habitats. Ecology 75, 2–16. 10.2307/1939377.

van Baalen M (1998). Coevolution of recovery ability and virulence. Proceedings of the Royal Society of London. Series B: Biological Sciences 265, 317–325. 10.1098/rspb.1998.0298.

van Baalen M, Rand DA (1998). The unit of selection in viscous populations and the evolution of altruism. Journal of theoretical biology 193, 631–648. 10.1006/jtbi.1998.0730.

van Baalen M, Sabelis MW (1995). The dynamics of multiple infection and the evolution of virulence. The American Naturalist 146, 881–910. 10.1086/590958.

Walsman JC, Duffy MA, Cáceres CE, Hall SR (2023). “Resistance is futile”: Weaker selection for resistance by abundant parasites increases prevalence and depresses host density. The American Naturalist 201, 864–879. 10.1086/724426.

Webster J, Gower C, Blair L (2004). Do hosts and parasites coevolve? Empirical support from the Schistosoma system. the american naturalist 164, S33–S51. 10.1086/424607.

