## Supplementary Material for "Double Trouble: Multiple infections and the coevolution of virulence and resistance in nested host-parasite populations" (from another version of this preprint: version 2, 2026-03-25)

### Appendix A : Supplementary informations

**Table 1** – Model parameters description

| Raw parameter | Biological Meaning | Unit | Scale |
|---|---|---|---|
| $\theta$ | Parasite investment in replication | None | Within-host |
| $\sigma$ | Host investment in resistance | None | Within-host |
| $b_0$ | Maximal host birth rate | individual.time$^{-1}$ | Between-host |
| $d$ | Host death rate | individual.time$^{-1}$ | Between-host |
| $\tau_0$ | Maximal recovery rate | individual.time$^{-1}$ | Between-host |
| $r_0$ | Maximal replication rate | individual.time$^{-1}$ | Within-host |
| $r_1$ | Strength of density dependance on parasite natality | None | Within-host |
| $\rho$ | Dispersal cost | None | Between-host |
| $m$ | Parasite dispersal rate | population.time$^{-1}$ | Within-host |
| $\mu$ | Parasite death rate | population.time$^{-1}$ | Within-host |
| $\gamma$ | Resistance rate | population.time$^{-1}$ | Within-host |
| Composite parameter | | | |
| $c = m(1-\rho)$ | Effective dispersal rate | individual.time$^{-1}$ | Between-host |
| $\beta(\theta, \sigma) = cX^*(\theta, \sigma)$ | Infectious contact rate | individual.time$^{-1}$ | Between-host |
| $\psi$ | Inverse of infectious period | time.individual$^{-1}$ | Between-host |
| $\phi_{mr}$ | Initial Fraction of the mutant when reinvaded by a resident | None | Within-host |
| $\phi_{rm}$ | Initial Fraction of the mutant when reinvading a resident patch | None | Within-host |
| $R$ | Relatedness | None | Within-host |
| $\bar{R} = 1 - R$ | Unrelatedness | None | Within-host |

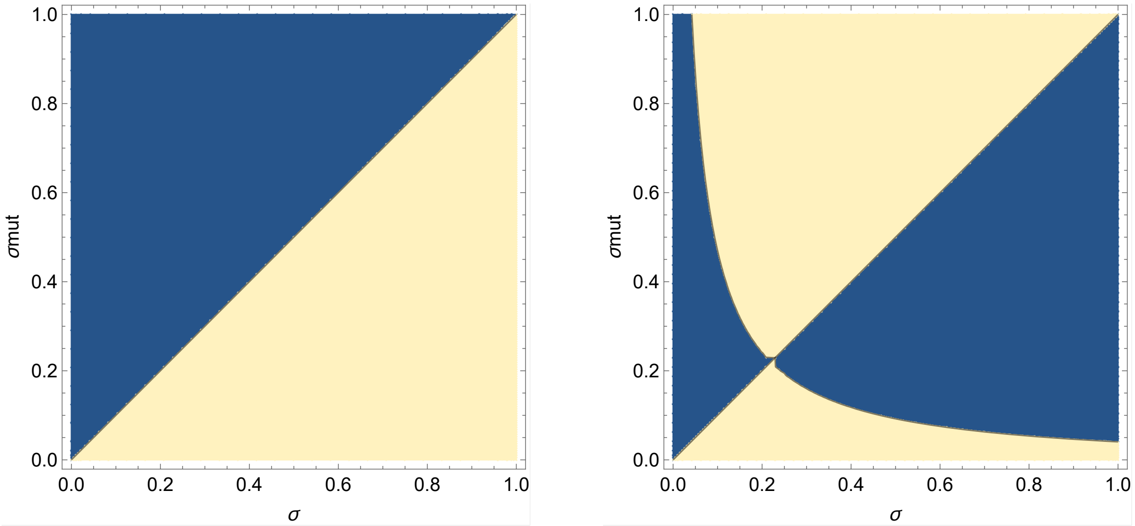


**(a)** **(b)**

**Figure 1** – Pairwise invasibility plots for the host evolving alone. White areas refer to cases where an invading parasite have a positive fitness $W_h > 1$, while blue areas refer to cases where a mutant parasite fades out. Fixed parameters were set to : :$b_0 = 0.8, \mu = 0.5, \alpha_0 = 1, r_0 = 10, \theta = 0.8, r_1 = 0.04, \rho = 0.9, \gamma_0 = 1, d_0 = 0.05, \tau_0 = 1, \delta = 3, \kappa = 1, d = 0.5$. (a) Moderately convex trade-off function between host fecundity and resistance ($\epsilon = 0.2$) (b) Strongly convex trade-off function between host fecundity and resistance ($\epsilon = 0.01$)

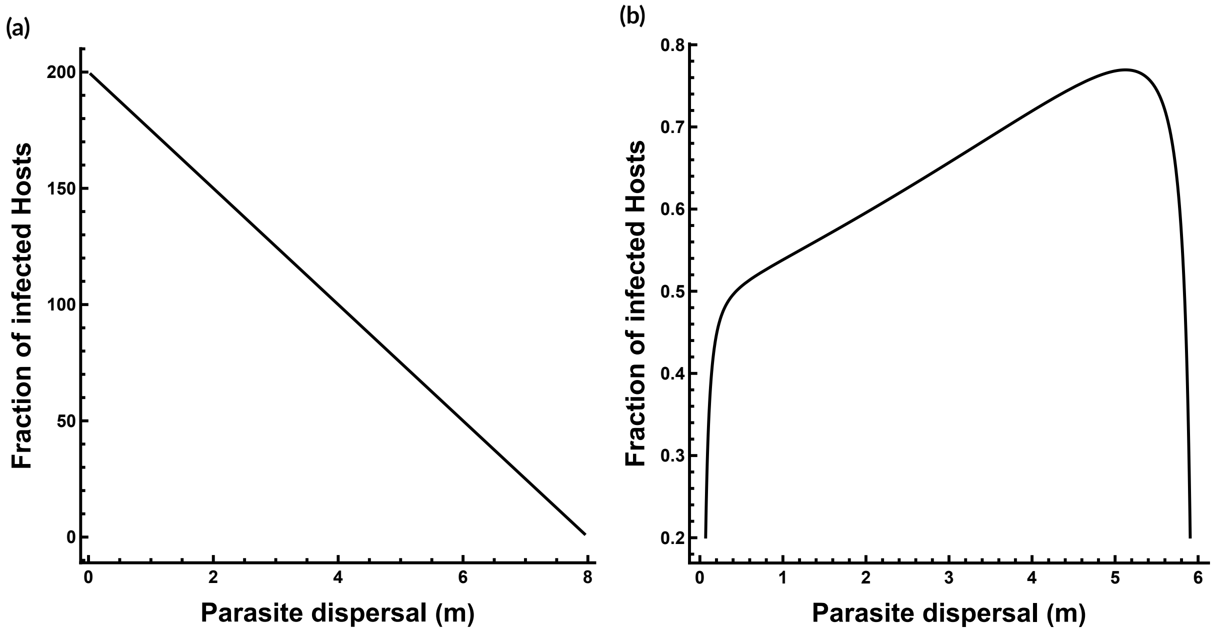


**Figure 2** – a) Fraction of infected host patches ($p_I^*$) as a function of parasite dispersal (m) b) Within-host equilibrium parasite load $X^*$, as a function of parasite dispersal (m) Fixed parameters were set to :$\theta = 0.9$, $\sigma = 0.5$, $\epsilon = 5$, $b_0 = 0.8$, $\mu = 0.5$, $\alpha_0 = 0.5$, $r_0 = 10$, , $r_1 = 0.04$, $\rho = 0.9$, $\gamma_0 = 1$, $d_0 = 0.05$, $\tau = 1$, $\delta = 5$, $\kappa = 1$, $\eta = 1$.

#### Appendix B : Host-driven parasite exintction

From system (3) in the main text, we can determine that the host is able to drive its parasite towards local extinction *iff* :

$$\frac{r_0\theta - (\mu + m)}{\gamma} \leq \sigma \leq 1 \tag{1}$$

It is then possible to constrain our parameters to satisfy the condition $r_0 - (\mu + m) > 0$, while allowing for the existence of a value $\sigma \in [0, 1]$ such that the within-host equilibrium density $X^*(\theta, \sigma) \leq 0$. This case is satisfied when $r_0\theta - (\mu + m) \leq \sigma \leq \frac{1}{\gamma}$.

In words, the ecological persistence of the parasite is always guaranteed in absence of host response, but there exists a level of resistance such that the host can drive its parasite to extinction.

In this scenario, studying the host evolution leads us to find that selection always drives resistance towards values that lead to parasite extinction. This singular strategy is however not an ESS in the strict sense, because once the singularity is reached, any mutant invader will be selectively neutral. This is an expected feature of our model, as we assumed that costs of resistance are triggered by immune up-regulation. As no maintenance costs are considered, the presence of the parasite is the only feature that creates selection upon host resistance.

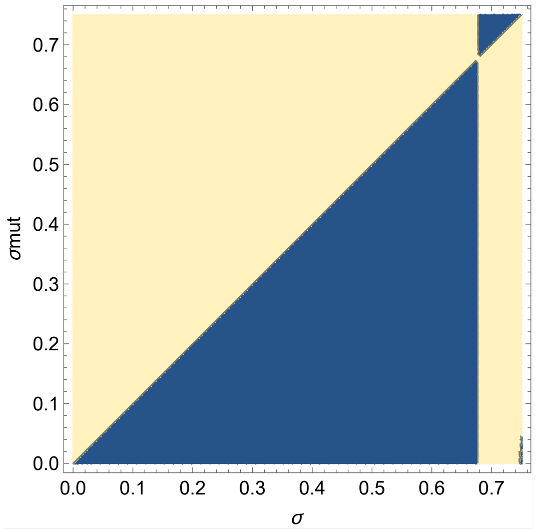


**Figure 3** – Pairwise invasibility plot for the host evolution with. Fixed parameter values for figures a-d were set to :$b_0 = 0.8, \mu = 0.5, \alpha_0 = 1, r_0 = 3, \theta = 0.8, r_1 = 0.04, \rho = 0.9, \gamma_0 = 1, d_0 = 0.05, \tau_0 = 1, \delta = 3, \kappa = 1, d = 0.5$

### Appendix C: Trade-off functions

In order to clarify the links between epidemiological and within-host parameters, we describe here the trade-off functions and the way transmission, virulence and recovery emerge from within-host traits.

We consider that both the host and the parasite have a finite amount of metabolic resources to allocate towards survival and/or reproduction. This allocation is determined by a (host or parasite) investment trait with values between 0 and 1. Our evolutionary analysis will investigate the evolution of these investment traits.

*Host fecundity and resistance.* Let $\sigma$ be the host investment in resistance. At the within-host scale, investment into resistance depletes parasite density at rate $\gamma\sigma$, where $\gamma$ is the maximum resistance level of a host. We assume that investment into resistance comes at the cost of a reduced fecundity, which, following Koffel et al. (2018), we express through the following trade-off function:

$$b(\sigma) = b_0\left(1 - \frac{\sigma}{\epsilon - (\epsilon - 1)\sigma}\right) \quad (2)$$

where $\epsilon > 0$ is a parmeter controlling the shape of the trade-off, as shown in figure (4). Note that we assume that the cost of resistance is expressed only in infected individuals, such that susceptible hosts always reproduce at rate $b(0) = b_0$.

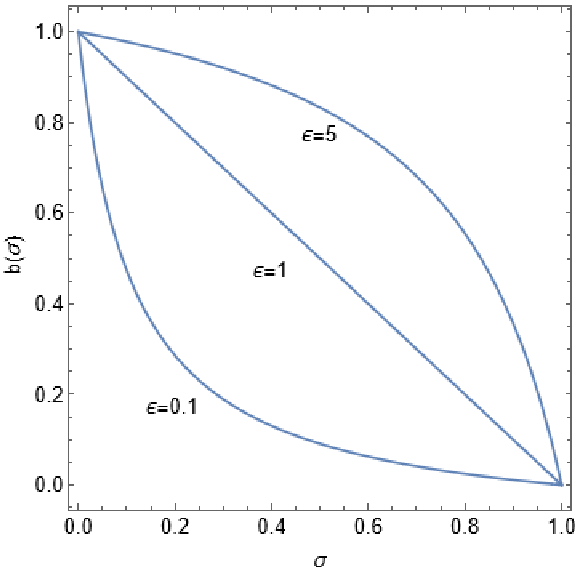


**Figure 4** – $b_0(\sigma)$ as a function of $\sigma$ for several values of $\epsilon$

*Parasite replication and load-dependant virulence.* Let $\theta$ be the parasite investment in replication. Similarly to host resistance, investment in parasite replication increases within-host parasite density at a rate $r_0\theta$, where $r_0$ is the maximal replication rate $r_0$ of a parasite. Our model assumes that investment in replication increases parasite load, which is positively correlated with both higher transmission and higher host mortality. Noting $X_{max}$ the maximum parasite load attainable by a parasite population, we define virulence as:

$$\alpha = f_\alpha(X^*) = \alpha_0 \frac{\frac{X^*}{X_{\alpha,max}}}{\delta - (\delta - 1)\frac{X^*}{X_{\alpha,max}}} \quad (3)$$

Here $\frac{X^*}{X_{\alpha,max}}$ is the relative equilibrium density of the parasite, with respect to its theoretical maximum defined by $X_{\alpha,max} = X^*(1, 0)$, and $\delta$ is a parameter controlling the shape of the trade-off. The use of the relative density $\frac{X^*}{X_{\alpha,max}}$ ensures that $f$ always remain bounded on $]0, 1[$

*Load dependant recovery.* We assume that recovery is a decreasing function of parasite load, that is, a host recovers faster when harbouring low densities of parasites. We assume:

$$\tau = f_\tau(X^*) = \tau_0 \frac{\left(1 - \frac{X^*}{X_{\tau,max}}\right)}{\kappa - (\kappa - 1)\left(1 - \frac{X^*}{X_{\tau,max}}\right)} \tag{4}$$

where $\frac{X^*}{X_{\tau,max}}$ is the relative equilibrium density of the parasite, with respect to the density corresponding to the fastest recovery rate, *i.e.* when $\theta = 0$ and $\sigma = 1$.

In order to keep biological consistency, we impose further constraints on dynamical processes of the interaction between the parasite and its host. We assume that the parasite needs a minimal investment in replication to ensure a positive growth rate, such that evolution towards avirulence is prevented. We also restrain our exploration of evolutionary dynamics to biologically relevant scenarii, considering only the cases where the functions $\tau(\theta, \sigma)$ and $\alpha(\theta, \sigma)$ stay positive.

#### Appendix D : Derivation of host fitness

Around the resident endemic equilibrium, the dynamics of the mutant is given by the following ODE system :

$$
\begin{aligned}
\frac{dp_{Sm}}{dt} &= [b_s p_{Sm} + b_{Im} p_{Sm}] p_0^* - \beta_m p_{Sm} p_I^* - d p_{Sm} + \tau_m p_{Im} \\
\frac{dp_{Im}}{dt} &= \beta_m p_{Sm} p_I^* - (d + \tau_m + \alpha_m) p_{Im}
\end{aligned} \tag{5}
$$

Invasion fitness is given by studying jacobian matrix associated to (5) (Hurford et al., 2010), which is :

$$
J(\sigma', \sigma) = \begin{pmatrix} b_s p_0 - \beta_m p_I - d_0 & b_I p_0 + \tau_m \\ \beta_m p_I & -(d_0 + \tau_m + \alpha_m) \end{pmatrix} \tag{6}
$$

We derive invasion fitness using matrix decomposition according to the next-generation theorem (Diekmann et al., 1990, Hurford et al., 2010), with :

$$
F = \begin{pmatrix} b_s p_0 & b_I p_0 \\ 0 & 0 \end{pmatrix} \tag{7}
$$

$$
V = \begin{pmatrix} \beta_m p_I + d_0 & -\tau_m \\ -\beta_m p_I & d0 + \tau_m + \alpha_m \end{pmatrix} \tag{8}
$$

Given that V is inversible, the spectral radius of $F.V^{-1}$ gives the invasion fitness. Setting $\psi = d + \alpha + \tau$, it leads after simplification to :

$$
W_h = \frac{b_0 \Psi_m + b_m \beta p_I^*}{d \Psi_m + (d + \alpha_m) \beta p_I^*} p_0^* \tag{9}
$$

#### Appendix E : Competition dynamics in a coinfected host

Let $X_m(t)$ and $X_r(t)$ be the densities of the mutant and resident strain in a co-infected host at time $t$. We have:

$$\frac{\mathrm{d}X_m}{\mathrm{d}t} =(r_0\theta_m - r_1X)X_m - (\mu + m + \gamma\sigma)X_m$$
$$\frac{\mathrm{d}X_r}{\mathrm{d}t} =(r_0\theta_r - r_1X)X_m - (\mu + m + \gamma\sigma)X_r$$

Summing these two equations yields the dynamics of the total density $Y(t) = X_m(t) + X_r(t)$

$$\frac{\mathrm{d}Y}{\mathrm{d}t} = r_0[\theta_m f(t) + \theta_r(1 - f(t)) - r_1 Y]Y - (\mu + m + \gamma\sigma)Y$$

where $f_m(t) = X_m(t)/Y(t)$ is the mutant frequency at time $t$. We can now write the dynamics of $f$ as:

$$\frac{\mathrm{d}f}{\mathrm{d}t} = \frac{1}{Y}\frac{\mathrm{d}X_m}{\mathrm{d}t} - \frac{\mathrm{d}Y}{\mathrm{d}t}\frac{X_m}{Y^2}$$

which simplifies into

$$\frac{\mathrm{d}f}{\mathrm{d}t} = f(1-f)r_0(\theta' - \theta).$$

Together with the initial condition $f(0) = \phi$, this ODE has an explicit solution:

$$f_m^{\phi}(t) = \frac{\phi}{\phi + (1-\phi)\exp^{-t\Delta s}}$$

which is equation (12) in the main text, with $\Delta s = r_0(\theta' - \theta)$.

Because we assume that local (within-host) dynamics are faster than the global (between-host) dynamics, we consider that the total parasite load reaches a quasi-equilibrium value $Y^{\phi}(t)$ on a fast time scale. Setting the right-hand-side of the dynamics of $Y(t)$ to zeo, we obtain

$$Y^{\phi}(t) = \frac{r_0[\theta_m f_m^{\phi}(t) + \theta_r(1 - f_m^{\phi}(t))] - (\mu + m + \gamma\sigma)}{r_1}.$$

Using the expressions for the equilibrium densities in singly infected hosts, we have

$$Y^{\phi}(t) = f_m^{\phi}(t)X_m^* + (1 - f_m^{\phi}(t))X_r^*.$$

The quasi-equilibrium parasite load in co-infected hosts is thus a weighted average of the densities in singly infected hosts, which change through time with the relative frequencies of the two strains.

#### Appendix F : From the parasite fitness function to the selection gradient

Our starting point is the following expression for the invasion fitness:

$$W_p = cp_S^* \left[ \frac{X_m^*}{\eta\beta p_I^* + \Psi_m} + \frac{\eta\beta p_I^*}{\eta\beta p_I^* + \Psi_m} \int_0^\infty F_m^{\phi_{mr}}(t)\mathrm{dt} \right] + \eta c p_1^* \int_0^\infty F_m^{\phi_{rm}}(t)\mathrm{dt} \tag{10}$$

Here :

- all quantities that depend on the mutant trait have a $m$ subscript.
- all quantities without subscripts solely depend on the resident trait.
- $\beta = cX^*$ and $\beta_m = cX_m^*$
- explicit dependence on the traits is dropped for simplicity.
- $X^*$ and $X_m^*$ refer to the equilibrium within-host parasite density in a singly infected host ($X_m^*$ : host infected by the mutant only; $X^*$ host infected by the resident).
- $F_m^{\phi}(t)$ is the number of mutants produced by doubly infected hosts at time t, with the superscript denoting the nature of reinvasion ($\phi_{rm}$ = reinvasion by the mutant , $\phi_{mr}$ = reinvasion by the resident). Note that $F_m^{\phi}(t)$ depends on both mutant and resident strains.

More precisely, we have :

$$F_m^{\phi}(t) = Y^{\phi}(t) f_m^{\phi}(t) \exp^{-\int_0^t \psi_\phi(s)ds} \tag{11}$$

Here $Y^{\phi}(t))$ is the quasi-equilibrium total density of parasite in doubly infected hosts (see Appendix E), $f_m^{\phi}(t)$ is the frequency of the mutant parasite at time $t$ in doubly infected hosts, and $\exp^{-\int_0^t \psi_\phi(s)ds}$ is a survival function that describes the probability that the infection has not ended at $t$.

Thus we can write the invasion fitness as $W_p = W_p^s + W_p^{\phi_{mr}} + W_p^{\phi_{rm}}$ with :

$$W_p^s = \text{single infections} = cp_S^* \frac{X_m^*}{\eta\beta p_I^* + \Psi_m} \tag{12a}$$

$$W_p^{\phi_{mr}} = \text{double infections } \phi_{mr} = cp_S^* \frac{\eta\beta p_I^*}{\eta\beta p_I^* + \Psi_m} \int_0^\infty F_m^{\phi_{mr}}(t)\mathrm{dt} \tag{12b}$$

$$W_p^{\phi_{rm}} = \text{double infections } \phi_{rm} = cp_1^* \int_0^\infty F_m^{\phi_{rm}}(t)\mathrm{dt} \tag{12c}$$

The selection gradient is given by the sum of the derivatives of these terms.

##### Single infections

The derivation of this term is straightforward and leads to :

$$\frac{\partial W_p^s}{\partial \theta_m} = \frac{cp_s^* \frac{\partial X_m^*}{\partial \theta_m}(\eta\beta p_I^* + \Psi_m) - cp_s^* X_m^* \frac{\partial \Psi_m}{\partial \theta_m}}{(\eta\beta p_I^* + \Psi_m)^2} \tag{13}$$

which can be simplified at neutrality, knowing that the resident $R_0$ is necessarily equal to one at its own equilibrium, it follows that $cp_S^* X^* = \Psi$. Using this simplification leads to :

$$\frac{\partial W_p^s}{\partial \theta_m}\Big|_{\theta_m=\theta} = \frac{\Psi}{\eta\beta p_I^* + \Psi} \left[ \frac{1}{X^*} \frac{\partial X_m^*}{\partial \theta_m}\Big|_{\theta_m=\theta} - \frac{1}{\eta\beta p_I^* + \Psi} \frac{\partial \Psi_m}{\partial \theta_m}\Big|_{\theta_m=\theta} \right] \tag{14}$$

**Double infections $\phi_{mr}$**

For coinfections resulting from the reinvasion by the resident strain of a host infected by the mutant, we obtain

$$\frac{\partial W_p^{\phi_{mr}}}{\partial \theta_m} = c p_s^* \Bigg[ \underbrace{-\frac{\eta\beta p_I^*}{(\eta\beta p_I^* + \Psi_m)^2}\frac{\partial \Psi_m}{\partial \theta_m}\int_0^\infty F_m^{\phi_{mr}}(t)\partial t}_{\text{A}} + \underbrace{\frac{\eta\beta p_I^*}{\eta\beta p_I^* + \Psi_m}\int_0^\infty \frac{\partial F_m^{\phi_{mr}}(t)}{\partial \theta_m}\,\mathrm{dt}}_{\text{B}} \Bigg] \tag{15}$$

The term A can be easily simplified using identities that hold at neutrality. Using :

$$Y^\phi(t)\Big|_{\theta_m=\theta} = X^*$$

$$f_m^\phi(t)\Big|_{\theta_m=\theta} = \phi$$

$$\exp^{-\int_0^t \psi_\phi(s)ds}\Big|_{\theta_m=\theta} = \exp^{-\Psi t}$$

we obtain after simplification

$$A = -\frac{\eta\beta p_I^*}{(\eta\beta p_I^* + \Psi_m)^2}\frac{\partial \Psi_m}{\partial \theta_m} X^* \phi_{mr}\frac{1}{\Psi} \tag{16}$$

The term B can be simplified using the appropriate identities that hold at neutrality :

$$\frac{\partial Y^\phi(t)}{\partial \theta_m}\Big|_{\theta_m=\theta} = \phi\frac{\partial X_m^*}{\partial \theta_m}\Big|_{\theta_m=\theta} \tag{17a}$$

$$\frac{\partial f^\phi(t)}{\partial \theta_m}\Big|_{\theta_m=\theta} = \frac{\partial \phi}{\partial \theta_m} + \phi(1-\phi)r_0 t \tag{17b}$$

$$\phi_{rm} = 1 - \phi_{mr} \tag{17c}$$

In addition, we note that

$$\begin{aligned}\frac{\partial F_m^{\phi_{mr}}(t)}{\partial \theta_m}\Big|_{\theta_m=\theta} &= \phi_m^{mr} e^{-\Psi t}\frac{\partial Y^{\phi_{mr}}(t)}{\partial \theta_m}\Big|_{\theta_m=\theta} + X^* e^{-\Psi t}\frac{\partial f^{\phi_{mr}}(t)}{\partial \theta_m}\Big|_{\theta_m=\theta} \\ &+ X^*\phi_m^{mr}\frac{\partial}{\partial \theta_m}\Big|_{\theta_m=\theta}\left(e^{-\int_0^t \Psi^{mr}(s)\,ds}\right).\end{aligned} \tag{18}$$

.

We also have

$$\frac{\partial}{\partial \theta_m}\left(e^{-\int_0^t \Psi^{mr}(s)\,ds}\right)\Big|_{\theta_m=\theta} = -e^{-\Psi t}\int_0^t \frac{\partial \Psi^{mr}(s)}{\partial \theta_m}\Big|_{\theta_m=\theta} ds \tag{19}$$

From the definition of $\Psi^{mr}$ we have, using the chain rule :

$$\frac{\partial \psi_\phi(s)}{\partial \theta_m}\Big|_{\theta_m=\theta} = (\alpha'(X^*) + \tau'(X^*))\,\frac{\partial X^{*\phi}(s)}{\partial \theta_m}\Big|_{\theta_m=\theta} \tag{20}$$

which reduces to :

$$\frac{\partial \psi_\phi(s)}{\partial \theta_m}\Big|_{\theta_m=\theta} = \phi\frac{\partial \Psi_m}{\partial \theta_m}\Big|_{\theta_m=\theta}. \tag{21}$$

Putting everything together gives us

$$\begin{aligned}\frac{\partial F_m^{\phi_{mr}}(t)}{\partial\theta_m} =&\phi_{mr}\frac{\partial X_m^*}{\partial\theta_m}f^{\phi_{mr}}(t)\exp^{-\int_0^t\psi_{\phi_{mr}}(s)ds}\partial t\\ &+X^{*\phi_{mr}}(t)\left[\frac{\partial\phi_{mr}}{\partial\theta_m}+\phi_{mr}(1-\phi_{mr})r_0t\right]\exp^{-\int_0^t\psi_{\phi_{mr}}(s)ds}\\ &+X^{*\phi_{mr}}(t)f^{\phi_{mr}}(t)\left[-\phi\frac{\partial\Psi_m}{\partial\theta_m}t\right]\exp^{-\int_0^t\psi_{\phi_{mr}}(s)ds}.\end{aligned}\tag{22}$$

which becomes, after evaluation at neutrality :

$$\begin{aligned}\frac{\partial F_m^{\phi_{mr}}(t)}{\partial\theta_m}\Big|_{\theta_m=\theta} =&\phi_{mr}^2\frac{\partial X_m^*}{\partial\theta_m}\Big|_{\theta_m=\theta}\exp^{-\Psi t}\\ &+X^*\left[\frac{\partial\phi_{mr}}{\partial\theta_m}\Big|_{\theta_m=\theta}+\phi_{mr}(1-\phi_{mr})r_0t\right]\exp^{-\Psi t}\\ &-X^*\phi_{mr}^2\frac{\partial\Psi_m}{\partial\theta_m}\Big|_{\theta_m=\theta}t\exp^{-\Psi t}.\end{aligned}\tag{23}$$

Then, integrating over $t$ gives :

$$\begin{aligned}\int_0^\infty\frac{\partial F_m^{\phi_{mr}}(t)}{\partial\theta_m}\Big|_{\theta_m=\theta}\partial t =&\phi_{mr}^2\frac{\partial X_m^*}{\partial\theta_m}\Big|_{\theta_m=\theta}\frac{1}{\Psi}\\ &+X^*\left[\frac{\partial\phi_{mr}}{\partial\theta_m}\Big|_{\theta_m=\theta}\frac{1}{\Psi}+\phi_{mr}(1-\phi_{mr})r_0\frac{1}{\Psi^2}\right]\\ &-X^*\phi_{mr}^2\frac{\partial\Psi_m}{\partial\theta_m}\Big|_{\theta_m=\theta}\frac{1}{\Psi^2}\end{aligned}\tag{24}$$

which leads, for the second term $B$ between brackets evaluated at neutrality, to

$$\begin{aligned}B =&\frac{\eta\beta p_I^*}{\eta\beta p_I^*+\Psi_m}\left[\phi_{mr}^2\frac{\partial X_m^*}{\partial\theta_m}\Big|_{\theta_m=\theta}\frac{1}{\Psi}\right.\\ &+X^*\left(\frac{\partial\phi_{mr}}{\partial\theta_m}\Big|_{\theta_m=\theta}\frac{1}{\Psi}+\phi_{mr}(1-\phi_{mr})r_0\frac{1}{\Psi^2}\right)\\ &\left.-X^*\phi_{mr}^2\frac{\partial\Psi_m}{\partial\theta_m}\Big|_{\theta_m=\theta}\frac{1}{\Psi^2}\right]\end{aligned}\tag{25}$$

Finally, using the expressions for $A$ and $B$, we obtain :

$$\begin{aligned}\frac{\partial W_p^{\phi_{mr}}}{\partial\theta_m}\Big|_{\theta_m=\theta} =&\frac{\eta\beta p_I^*}{(\eta\beta p_I^*+\Psi)}\left[-\frac{1}{(\eta\beta p_I^*X^*+\Psi)}\frac{\partial\Psi_m}{\partial\theta_m}\phi_{mr}+\phi_{mr}^2\frac{1}{X^*}\frac{\partial X_m^*}{\partial\theta_m}\Big|_{\theta_m=\theta}\right.\\ &\left.+\frac{\partial\phi_{mr}}{\partial\theta_m}\Big|_{\theta_m=\theta}+\phi_{mr}(1-\phi_{mr})r_0\frac{1}{\Psi}-\phi_{mr}^2\frac{\partial\Psi_m}{\partial\theta_m}\Big|_{\theta_m=\theta}\frac{1}{\Psi}\right]\end{aligned}\tag{26}$$

**Double infections $\phi_{rm}$**

The third term can be obtained by simply swapping indices $r$ and $m$, which yields :

$$\frac{\partial W_p^{\phi_{rm}}}{\partial \theta_m}\Big|_{\theta_m=\theta} = \eta\beta p_1^* \frac{1}{\Psi}\Big[\phi_{rm}^2 \frac{1}{X^*}\frac{\partial X_m^*}{\partial \theta_m}\Big|_{\theta_m=\theta} + \frac{\partial \phi_{rm}}{\partial \theta_m}\Big|_{\theta_m=\theta} + \phi_{rm}(1-\phi_{rm}) r_0 \frac{1}{\Psi} - \phi_{rm}^2 \frac{\partial \Psi_m}{\partial \theta_m}\Big|_{\theta_m=\theta}\frac{1}{\Psi}\Big] \tag{27}$$

Noting that :

$$p_1^* = \frac{\eta\beta p_l^* p_S^*}{\eta\beta p_l^* + \Psi} \tag{28}$$

We obtain :

$$\frac{\partial W_p^{\phi_{rm}}}{\partial \theta_m}\Big|_{\theta_m=\theta} = \frac{\eta\beta p_l^*}{\eta\beta p_l^* + \Psi}\Big[\phi_{rm}^2 \frac{1}{X^*}\frac{\partial X_m^*}{\partial \theta_m}\Big|_{\theta_m=\theta} + \frac{\partial \phi_{rm}}{\partial \theta_m}\Big|_{\theta_m=\theta} + \phi_{rm}(1-\phi_{rm}) r_0 \frac{1}{\Psi} - \phi_{rm}^2 \frac{\partial \Psi_m}{\partial \theta_m}\Big|_{\theta_m=\theta}\frac{1}{\Psi}\Big] \tag{29}$$

**Fitness gradient**

If we sum the three derivatives, we obtain

$$\begin{aligned}\frac{\partial W_p}{\partial \theta_m}\Big|_{\theta_m=\theta} =& \frac{\Psi}{\eta\beta p_l^* + \Psi}\Big[\frac{1}{X^*}\frac{\partial X_m^*}{\partial \theta_m}\Big|_{\theta_m=\theta} - \frac{1}{\eta\beta p_l^* + \Psi}\frac{\partial \Psi_m}{\partial \theta_m}\Big|_{\theta_m=\theta}\Big] \\ &+ \frac{\eta\beta p_l^*}{(\eta\beta p_l^* + \Psi)}\Big[-\frac{1}{(\eta\beta p_l^* + \Psi)}\frac{\partial \Psi_m}{\partial \theta_m}\phi_{mr}\Big] \\ &+ \frac{\eta\beta p_l^*}{(\eta\beta p_l^* + \Psi)}\Big[\phi_{mr}^2 \frac{1}{X^*}\frac{\partial X_m^*}{\partial \theta_m}\Big|_{\theta_m=\theta} \frac{\partial \phi_{mr}}{\partial \theta_m}\Big|_{\theta_m=\theta} + \phi_{mr}(1-\phi_{mr}) r_0 \frac{1}{\Psi} - \phi_{mr}^2 \frac{\partial \Psi_m}{\partial \theta_m}\Big|_{\theta_m=\theta}\frac{1}{\Psi}\Big] \\ &+ \frac{\eta\beta p_l^*}{\eta\beta p_l^* + \Psi}\Big[\phi_{rm}^2 \frac{1}{X^*}\frac{\partial X_m^*}{\partial \theta_m}\Big|_{\theta_m=\theta} \frac{\partial \phi_{rm}}{\partial \theta_m}\Big|_{\theta_m=\theta} + \phi_{rm}(1-\phi_{rm}) r_0 \frac{1}{\Psi} - \phi_{rm}^2 \frac{\partial \Psi_m}{\partial \theta_m}\Big|_{\theta_m=\theta}\frac{1}{\Psi}\Big]\end{aligned} \tag{30}$$

Finally, we use the fact that at neutrality we have

$$\begin{aligned}\frac{\partial \phi_{mr}}{\partial \theta_m}\Big|_{\theta_m=\theta} &= -\frac{\partial \phi_{rm}}{\partial \theta_m}\Big|_{\theta_m=\theta} \\ \phi_{rm} + \phi_{mr} &= 1\end{aligned} \tag{31}$$

and get a simpler expression for the selection gradient

$$\begin{aligned}\frac{\partial W_p}{\partial \theta_m}\Big|_{\theta_m=\theta} =& \frac{\Psi}{\eta\beta p_l^* + \Psi}\Big[\frac{1}{X^*}\frac{\partial X_m^*}{\partial \theta_m}\Big|_{\theta_m=\theta} - \frac{1}{\eta\beta p_l^* + \Psi}\frac{\partial \Psi_m}{\partial \theta_m}\Big|_{\theta_m=\theta}\Big] \\ &+ \frac{\eta\beta p_l^*}{(\eta\beta p_l^* + \Psi)}\Big[-\frac{1}{(\eta\beta p_l^* + \Psi)}\frac{\partial \Psi_m}{\partial \theta_m}\phi_{mr} \\ &+ (\phi_{mr}^2 + \phi_{rm}^2)\Big(\frac{1}{X^*}\frac{\partial X_m^*}{\partial \theta_m}\Big|_{\theta_m=\theta} - \frac{1}{\Psi}\frac{\partial \Psi_m}{\partial \theta_m}\Big|_{\theta_m=\theta}\Big) + 2\phi_{mr}\phi_{rm} r_0 \frac{1}{\Psi}\Big]\end{aligned} \tag{32}$$

which after rearrangement, and using $\beta_m = cX_m^*$, leads to the form used in the main text :

$$\frac{\partial W_p}{\partial \theta_m}\Big|_{\theta_m=\theta} = \frac{\Psi}{\eta\beta p_I^* + \Psi}\left[\frac{\partial}{\partial\theta_m}\left(\ln\frac{\beta_m}{\Psi_m + \eta\beta p_I^*}\right)\Big|_{\theta_m=\theta}\right] + \frac{\eta\beta p_I^*}{\eta\beta p_I^* + \Psi}\left[-\frac{1}{\eta\beta p_I^* + \Psi}\phi_{mr}\frac{\partial\Psi_m}{\partial\theta_m}\Big|_{\theta_m=\theta} + \left(\frac{\partial}{\partial\theta_m}\left(\ln\frac{\beta_m}{\Psi_m}\right)\Big|_{\theta_m=\theta}\right)(\phi_{mr}^2 + \phi_{rm}^2) + 2\phi_{mr}\phi_{rm}r_0\frac{1}{\Psi}\right]$$

#### Unravelling relatedness

Eq.(32) can be further interpreted by noticing that its third term can be understood in terms of relatedness between parasite genotypes in coinfected hosts. We compute relatedness in the metapopulation following the methods in (Rousset and Billiard, 2000), (Jansen and Vitalis, 2007)(Appendix 3).

Consider a metapopulation at its equilibrium, populated by two strains selectively neutral labelled $m$ and $r$. The total number of parasite in the metapopulation is $X^* p_I N$ where $p_I$ is the fraction of infected individuals (with no regard on coinfection and number of invasions) and $N$ an arbitrary number of hosts. Among the whole set of parasites, let $\omega$ the fraction of $m$ strain, and $(1-\omega)$ the fraction of $r$ strain. As the dynamics of the two-strains system equilibrates, so does the distribution of individuals among patches. Therefore if $p_1^*$ is the fraction of infected individuals invaded once, a fraction $\omega$ of these individuals has been invaded by the strain $m$, and a fraction $(1-\omega)$ by the strain $r$. With the same reasonning, if $p_2^*$ is the fraction of individuals invaded twice, then a fraction $\omega^2$ of those patches has been invaded by the strain $m$, and a fraction $(1-\omega)^2$ by the strain $r$. The fraction of patches that exhibit coinfection is therefore $1 - w^2 - (1-w)^2 = 2w(1-w)$.

To derive relatedness in our model, we focus on the subset of infected individuals that experienced two parasite invasions, possibly by the same strain (*i.e.* the fraction $p_2^*$).

Thus, the probability to sample to identical individual $m$ in the whole (meta)population of hosts is

$$p_m^b = \omega^2, \tag{33}$$

and the probability to sample to identical individuals $r$ is therefore

$$p_b^r = (1-\omega)^2 \tag{34}$$

.

Similarly, the probability to sample two identical individuals $m$ in a given unique host is :

$$p_w^m = \frac{\omega p_1^* + \omega^2 p_2^* + \omega(1-\omega)p_2^*(\phi_{rm}^2 + \phi_{mr}^2)}{p_I^*}$$

and the probability to sample identical individuals $r$ is

$$p_w^r = \frac{(1-\omega)p_1^* + (1-\omega)^2 p_2^* + \omega(1-\omega)p_2^*(\phi_{rm}^2 + \phi_{mr}^2)}{p_I^*}$$

The within-host relatedness, relative to the (meta)population of hosts is computed as :

$$R = \frac{p_w^m + p_w^r - p_b^r - p_b^m}{1 - p_b^m - p_b^r} = (\phi_{rm}^2 + \phi_{mr}^2) \tag{35}$$

and the unrelatedness is simply :

$$\bar{R} = 1 - R = 2\phi_{rm}\phi_{mr} \tag{36}$$

Relatedness then naturally emerges from eq.(33), which gives :

$$\begin{aligned} \frac{\partial W_p}{\partial \theta_m}\Big|_{\theta_m=\theta} =& \frac{\Psi}{\eta\beta p_l^* + \Psi}\left[\frac{\partial}{\partial \theta_m}\left(\ln \frac{\beta_m}{\Psi_m + \eta\beta p_l^*}\right)\Big|_{\theta_m=\theta}\right] \\ &+ \frac{\eta\beta p_l^*}{\eta\beta p_l^* + \Psi}\left[-\frac{1}{\eta\beta p_l^* + \Psi}\phi_{mr}\frac{\partial \Psi_m}{\partial \theta_m}\Big|_{\theta_m=\theta}\right. \\ &\left. + R\left(\frac{\partial}{\partial \theta_m}\left(\ln \frac{\beta_m}{\Psi_m}\right)\Big|_{\theta_m=\theta}\right) + \bar{R} r_0 \frac{1}{\Psi}\right] \end{aligned} \tag{37}$$

,

which is the expression used in the main text.

#### Appendix G : From coinfections to superinfection

We detail here the derivation of the invasion fitness of a mutant parasite in the case of superinfection, starting from our initial coinfection formulation (eq. 20 in the main text). Our starting point is thus the invasion fitness of a mutant parasite assuming multiple infection can occur and that the parasite strains transiently coexists in an infected host :

$$W_p = p_S^* \left[ \frac{\beta_m}{\eta \beta p_I^* + \Psi_m} + \frac{\eta_r \beta p_I^*}{\eta_r \beta p_I^* + \Psi_m} \int_0^\infty F_m^{\phi_{mr}}(t) \mathrm{d}t \right] + \eta_m p_1^* \int_0^\infty F_m^{\phi_{rm}}(t) \mathrm{d}t \tag{38}$$

In order to comply with the superinfection hypothesis, we now assume that following a secondary infection, the invading strain instantanesouly exclude the resident one with probability $\eta$, or fades out. That is, $\eta_r$ (resp. $\eta_m$) is the probability that the resident (resp. mutant) strain displaces the mutant (resp. resident) upon reinvasion. In this case, the coexistence of strains in a host is precluded.

With that consideration in mind, it follows that the initial frequency of a mutant strain after being successfully reinvaded by the resident strain is $\phi_{mr} = 0$. Conversely, the initial frequency of the mutant strain after it reinvades the resident is $\phi_{rm} = 1$.

Thus, the number of mutant propagules profuced after being reinvaded by the resident is straightforwardly

$$F_m^{\phi_{mr}}(t) = 0 \tag{39}$$

Because the strains cannot coexist in a host, it is not necessary anymore to restrict the number of infection, and the mutant strain can reinvade any host infected by the resident strain, without limitations. In addition, after a successful reinvasion has occured, the resident strain is able to reinvade again. Consequently, the number of propagules produced after the mutant reinvades the resident becomes

$$F_m^{\phi_{rm}}(t) = cX_m^* \int_0^\infty \exp^{-(\Psi + \beta_r \eta_r p_I^*)t} \, dt = \frac{\beta_m}{\Psi + \beta_r \eta_r p_I^*} \tag{40}$$

where $\int_0^\infty \exp^{-(\Psi+\beta_r\eta_r p_I^*)t} \, dt$ denotes the probability that the infection ends at time $t$. We can now consider the set of fates a mutant invader can experience.

If the mutant lands in a susceptible host $p_s^*$, it will therefore produce $\beta_m$ dispersers per unit of time until the host recovers, dies or is reinvaded by the resident strain. The reinvasion by the resident occurs at rate $\beta_r p_I^* \eta_r$, and the infection ends at a rate $\Psi_m$.

If the mutant lands in an infected host $p_I^*$, it will successfully establish with probability $\eta_m$. Following the reinvasion, it will produce $\beta_m$ propagules per unit of time until the infection ends or the resident strain invades the host again.

Putting the two scenarii together gives the invasion for a rare mutant in the case of superinfections :

$$W_p^s = \frac{\beta_m \eta_m p_I^*}{\Psi_m + \beta_r \eta_r p_I^*} + \frac{\beta_m p_S^*}{\Psi_m + \beta_r \eta_r p_I^*} \tag{41}$$

which, in the absence of the shared control of epidemiological traits, is equivalent to eq.(A3) in (Kada and Lion, 2015). Note that because $\beta = cX^*$ still depends on the within-host equilibrium load $X^*$, the rate at which the mutant replace the resident strain depends on its within-host competitive ability, and will still alter the duration of infection. The derivative of eqn.(41) with respect to $\theta_m$ leads to the equation (22) in the main text.

#### References

Diekmann O, Heesterbeek JAP, Metz JA (1990). *On the definition and the computation of the basic reproduction ratio R 0 in models for infectious diseases in heterogeneous populations*. Journal of mathematical biology **28**, 365–382. https://doi.org/10.1007/BF00178324.

Hurford A, Cownden D, Day T (2010). *Next-generation tools for evolutionary invasion analyses*. Journal of the Royal Society Interface **7**, 561–571. https://doi.org/10.1098/rsif.2009.0448.

Jansen VA, Vitalis R (2007). *THE EVOLUTION OF DISPERSAL IN A LEVINS'TYPE METAPOPULATION MODEL*. Evolution **61**, 2386–2397. https://doi.org/10.1111/j.1558-5646.2007.00201.x.

Kada S, Lion S (2015). *Superinfection and the coevolution of parasite virulence and host recovery*. Journal of evolutionary Biology **28**, 2285–2299. https://doi.org/10.1111/jeb.12753.

Koffel T, Daufresne T, Massol F, Klausmeier CA (2018). *Plant strategies along resource gradients*. The American Naturalist **192**, 360–378. https://doi.org/10.1086/698600.

Rousset, Billiard (2000). *A theoretical basis for measures of kin selection in subdivided populations: finite populations and localized dispersal*. Journal of Evolutionary Biology **13**, 814–825. https://doi.org/https://doi.org/10.1046/j.1420-9101.2000.00219.x.
